# Rearrangement of T Cell Genome Architecture Regulates GVHD

**DOI:** 10.1101/2021.01.23.427857

**Authors:** Yaping Sun, Gabrielle A. Dotson, Lindsey A. Muir, Scott Ronquist, Katherine Oravecz-Wilson, Daniel Peltier, Keisuke Seike, Lu Li, Walter Meixner, Indika Rajapakse, Pavan Reddy

**Affiliations:** Department of Internal Medicine, Division of Hematology and Oncology, University of Michigan Rogel Cancer Center, Ann Arbor, MI, USA; Department of Computational Medicine and Bioinformatics, University of Michigan, Ann Arbor, MI 48109 USA; Department of Mathematics, University of Michigan, Ann Arbor, MI, USA

## Abstract

The cohesin complex modulates gene expression and cellular functions by shaping three-dimensional (3D) organization of chromatin. WAPL, cohesin’s DNA release factor, regulates 3D chromatin architecture. The 3D genome structure and its relevance to mature T cell functions *in vivo* is not well understood. We show that *in vivo* lymphopenic expansion, and allo-antigen driven proliferation, alters the 3D structure and function of the genome in mature T cells. Conditional deletion of *Wapl* in T cells reduced long-range genomic interactions, altered chromatin A/B compartments and interactions within topologically associating domains (TADs) of the chromatin in T cells at baseline. Comparison of chromatin structure in normal and WAPL-deficient T cells after lymphopenic and allo-antigen driven stimulation revealed reduced loop extensions with changes in cell cycling genes. WAPL-mediated changes in 3D architecture of chromatin regulated activation, cycling and proliferation of T cells *in vitro* and *in vivo*. Finally, WAPL-deficient T cells demonstrated reduced severity of graft-versus-host disease (GVHD) following experimental allogeneic hematopoietic stem cell transplantation. These data collectively characterize 3D genomic architecture of T cells *in vivo* and demonstrate biological and clinical implications for its disruption by cohesin release factor WAPL.

## Introduction

The three-dimensional (3D) architecture of the genome includes coiling of genomic DNA around histone proteins to form the chromatin fiber, which folds into higher-order structures such as loops, domains, compartments, and chromosomes (Bonev and Cavalli, 2016; Finn and Misteli, 2019; Misteli, 2020; Rajapakse et al., 2011; Rajapakse and Groudine, 2011). High resolution chromatin conformation capture experiments reveal that the 3D spatial architecture of the chromatin at various scales is conserved, reproducible at the cellular level, and regulates gene expression (Cremer and Cremer, 2019, 2001; Dekker and Mirny, 2016; Dekker et al., 2002). It is increasingly appreciated that higher spatial organization can have specific alterations during mammalian development and in some pathologies including cancers and infections (Finn and Misteli, 2019; Waldman, 2020). However, whether *a priori* disruption of this 3D organization alters *in vivo* cellular functions and disease processes remains poorly understood.

The multi-unit cohesin ring complex plays a critical role in 3D genomic organization and in cell division. It consists of SMC1/3, SCC1 (RAD21), and STAG subunits that are loaded by the SCC2/SCC4 complex onto genomic DNA and establish the cohesin ring structure (Cuadrado and Losada, 2020; Nasmyth and Haering, 2009; Piché et al., 2019; Remeseiro et al., 2013). Cohesin dependency has been demonstrated by depletion of various cohesin units (Piché et al., 2019). Cohesin release from chromatin is driven by WAPL, which opens an exit site at the interface of the SMC3/SCC1 subunits of the cohesin ring (Haarhuis et al., 2013, 2017; Silva et al., 2020). Prior studies have elegantly demonstrated that the absence of WAPL reduces cohesin turnover, alters chromatin loop extensions, and leads to defects in interphase chromosome organization (Busslinger et al., 2017; Haarhuis et al., 2013, 2017; Hill et al., 2020; Silva et al., 2020; Tedeschi et al., 2013).

WAPL is essential during mammalian embryonic development (Tedeschi et al., 2013). However, the role of WAPL-dependent chromatin alterations on *in vivo* functions after embryonal development or in mature T cells is not known. The 3D chromatin landscape has been recently described in T cell development, in T cell lines and following *in vitro* stimulation (Hu et al., 2018; Isoda et al., 2017; Misteli, 2020; Rawlings et al., 2011; Robson et al., 2017; Yang et al., 2020; Burren et al., 2017; Bediaga et al., 2021). It remains unclear however whether the specific changes in the 3D chromatin landscape influence or emerge from T cell development. Furthermore, the functional chromatin landscape in mature T cells and their alterations following *in vivo* activation remain unknown. The role of cohesin ring formation in mature T cell development, its function *in vivo*, and its disruption by the absence of WAPL in mature T cell immunity are also not fully understood.

Mature T cells can cause graft-versus-host disease (GVHD) following allogeneic hematopoietic cell transplantation (HCT), a potentially curative therapy against many hematological malignant and non-malignant diseases (Blazar et al., 2020; Wu and Reddy, 2017; Zeiser and Blazar, 2017). GVHD has precluded widespread utilization of this effective therapy. The chromatin landscape of allogeneic T cells, and whether the disruption of this landscape can regulate the severity of T cell-mediated GVHD remains unexplored.

Herein, we utilized genome-wide chromosome conformation capture (Hi-C) and RNA-sequencing to describe the 3D chromatin architecture of mature T cells *in vivo*, specifically at baseline (unstimulated, naive, pre-transplant) and following non-antigen stimulated lymphopenia induced proliferation (syngeneic) and allo-antigen driven (allogeneic) stimulation (Maeda et al., 2007) (Figure S1). We evaluated 3D chromatin architecture at the chromosome, TAD, and sub-TAD levels to capture both global and local trends of T cell genome structure (Figure 2A). We generated T cell-specific WAPL-deficient mice and demonstrated that WAPL regulates the 3D chromatin structure, function, and *in vivo* biological response of T cells such as GVHD. These data collectively demonstrate that *a priori* disruption of chromatin structure regulates mature T cell function.

## Results

### Characterization of mature naïve T cell genome architecture following *in vivo* stimulation

We first determined the genome architecture of mature naïve T cells at baseline and following *in vivo* lymphopenic and antigen-driven stimulation. To mimic clinically-relevant *in vivo* stimulation, we compared naïve T cells before and after experimental syngeneic and allogeneic transplantation. To this end, we utilized MHC-disparate B6 into a BALB/c model of transplantation (Reddy et al., 2008). CD62L^+^ naive donor T cells were harvested from the splenocytes of B6 donors and transplanted into congenic syngeneic B6 or allogeneic BALB/c recipients (see Methods). Recipient animals were sacrificed seven days after transplantation and their splenic T cells were isolated using congenic markers and analyzed for genomic architecture. While we generated Hi-C contact maps to profile genome-wide chromatin interactions (Rao et al., 2014) for harvested T cells, variability in the number of mappable Hi-C reads among unstimulated and stimulated T cell settings (Table S1) precluded us from reliably making direct comparisons between the genome architecture of naive, syngeneic, and allogeneic T cells. From the RNA-seq data, however, we did observe significant differences in gene expression before and after *in vivo* stimulation, most notably in chromosomes 7 and 11 (Figure S2).

### Generation of T cell conditional WAPL-deficient mice

Because T cell function changed following stimulation, we next sought to evaluate which conformational features in the genome are critical for T cell function. Cohesin promotes chromatin looping while WAPL is important for the release of cohesin from chromatin. WAPL is essential for embryonal development and its deficiency has been shown to cause defects in chromatin structure (Tedeschi et al., 2013). Specifically, WAPL has been shown to restrict chromosome loop extension (Busslinger et al., 2017; Haarhuis et al., 2017; Haarhuis and Rowland, 2017; Hill et al., 2020). In addition, we previously demonstrated that mir142 regulates mature T cell pro-liferation, targets WAPL expression in T cells and that it may modulate T cell activation (Sun et al., 2013). Therefore, we determined whether WAPL regulates 3D chromatin architecture and the function of mature T cells.

Because WAPL is critical for embryonal development, we generated a T cell conditional *Wapl* knock-out (KO) mouse using CRISPR-Cas9 and CD4-CRE systems (Ran et al., 2015; Tabebordbar et al., 2016). The *Wapl* locus on chromosome 14q has 18 exons and one noncoding exon (Figure S3A). We designed two sgRNAs specific to exon 2 of the *Wapl* gene to generate a double strand break in exon 2 (Figure 1A and Figure S3B). Our first line of mice carried an insert of two sgRNAs targeting exon 2 in *Wapl* (Figure 1A and Figure S3C). The second and third lines carried Rosa26-floxed STOP-Cas9 knock-in on B6J (The Jackson Laboratory, Stock No:026175) or were CD4-CRE transgenic mice (The Jackson Laboratory, Stock No: 017336) (Figure S3C). Triple crosses were screened for sgRNA insert showing the positive 457 bp band (sgRNA-Wapl) (Figure 1B), CRE positive as a 100 bp band, loxP-SV40pA x3-loxP-CAS9-GFP (LSL) cleavage activity demonstrating 1,123 bp for WT LSL, and a 285 bp band for cleaved LSL after CRE recombination processing. The conditional KO mice developed normally. The *Wapl* KO T cells were verified for GFP expression (Figure 1B). The higher levels of LSL cleavage and GFP expression assured successful depletion of WAPL (Figure 1B, lane 4). We further confirmed WAPL protein depletion through Western blotting (Figure 1C) in T cells isolated from multiple knockout pups. Finally, to further confirm efficient deletion, we performed RNA-seq on the T cells sorted from these mice and compared them with littermate WT T cells, which demonstrated efficient loss of exon 2 in the *Wapl* gene (Figure 1D).

**Fig. 1:**
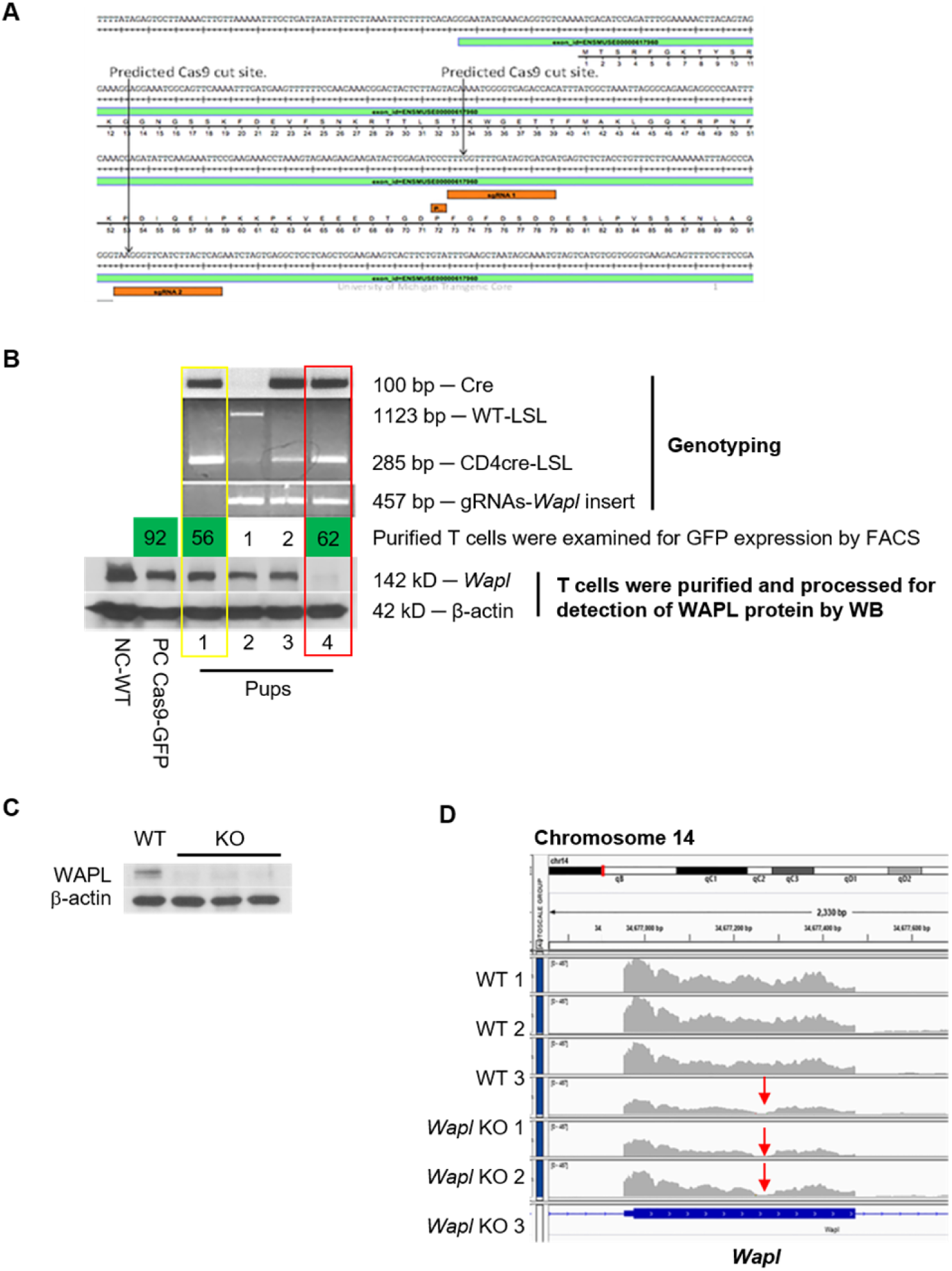
Generating conditional *Wapl* knockout T cells utilizing CRISPR-CAS9 system. (A) Map of two sgRNAs targeting exon 2 of the *Wapl* gene on Chromosome 14. (B) Genotyping and confirmation of *Wapl* knockout. Mice tail DNA was processed for PCR genotyping to confirm expression of CRE and sgRNA-*Wapl* insert and Cas9-stop signal expression or cleavage. GFP expression was determined by flow cytometry and WAPL protein deletion was determined by Western Blotting. (C) Confirmation of *Wapl* KO in T cells on multiple pups according to the screen procedures in (B) by Western Blotting. (D) RNA-seq confirmation across biological triplicates of sgRNA-mediated deletion of exon 2 in *Wapl*.

### WAPL regulates T cell genome architecture

Because WAPL is known to regulate genome architecture and genome architecture entrains transcription (Kueng et al., 2006; Tedeschi et al., 2013; Wutz et al., 2017), we next evaluated the impact of WAPL deficiency on genome architecture and gene expression of unstimulated naïve T cells. To analyze whole genome architecture (structure) and gene expression (function), we integrated Hi-C and RNA-seq data (see Methods). We performed RNA-seq on T cells harvested from naïve B6 and those harvested on day+7 from transplanted syngeneic and allogeneic B6 recipients (see Methods).

We evaluated the statistical dissimilarity between wildtype (WT) and knockout (KO) Hi-C contact matrices at the chromosome level in each setting by employing the Larntz-Perlman (LP) procedure (see Methods) which revealed regions critically impacted by *Wapl* knockout. WT and KO matrices were significantly dissimilar across all chromosomes (p *<* .001). Though statistically different, WT and KO chromosomes in unstimulated naïve T cells had a lesser number of differential regions as compared with chromosomes in syngeneic and allogeneic T cells (Figure 2B). This conservation of structure is consistent with the notion that unstimulated naïve T cells are in a quiescent-like state and therefore less impacted by the loss of WAPL. Further, differential regions in the naive setting appear more concentrated between ends of chromosomes – suggesting that long-range interactions may be critical to maintaining structural integrity in unstimulated T cells.

**Fig. 2:**
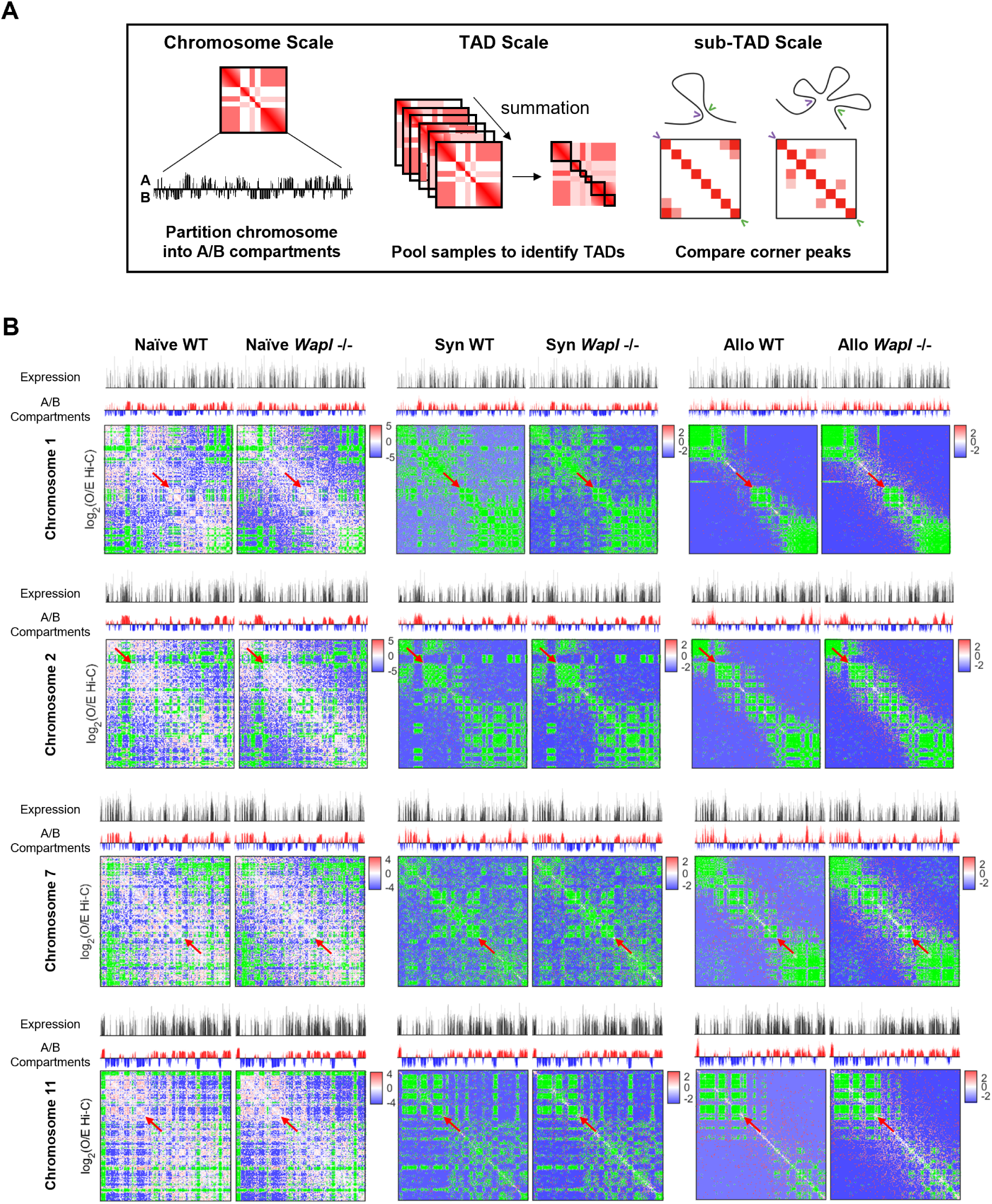
Genome-wide effects of WAPL knockout in unstimulated naïve T cells and after syngeneic/allogeneic transplantation. (A) Illustration of hierarchical Hi-C analysis workflow. (B) Chromosome-level features and contact maps for chromosomes 1 and 2 from wildtype and *Wapl* knockout T cells. Gene expression (top track) is shown as a vector of log_2_(TPM) values binned at 100kb resolution and chromatin accessibility (bottom track) is shown as signed values from the Fiedler vector of the Hi-C contact map where positive values (red) denote A compartments and negative values (blue) denote B compartments. ICE and O/E normalized Hi-C contact maps are shown at 100-kb resolution and log-scale. Matrix dissimilarity between WT and KO conditions was detected by the Larntz-Perlman procedure (see Methods) and genomic regions with dissimilarity in the 99th percentile are shaded in green. These areas indicate the largest perturbations to the chromatin architecture upon *Wapl* knockout.

We highlight regions in the 99th percentile of significant changes across chromosomes 1, 2, 7, and 11 in Figure 2B, due to their abundance of differential expression and compartmentalization (Figures S4 and S5). Across all naïve chromosomes, 15-25% of genes within these LP-dissimilar regions are significantly differentially expressed (p *<* 0.05) compared to 8.5-13% across LP-dissimilar regions of syngeneic chromosomes and 3-7% across LP-dissimilar regions of allogeneic chromosomes. Interestingly, there are differential regions in common between the syngeneic and allogeneic settings that are not detected in the naive setting (red arrows in Figure 2B), indicating stimulation-specific genome architecture.

We then bi-partitioned chromosomes into individual stretches of accessible (active) and inaccessible (inactive) chromatin, termed A (euchromatin) and B (heterochromatin) compartments, respectively (Lieberman-Aiden et al., 2009). We demarcated these regions using the signed values of the Fiedler vector which measures underlying chromatin accessibility (Chen et al., 2015). The positive values of the Fiedler vector reflect compartment A and negative values reflect compartment B. We observed A/B compartment switch events mediated by the loss of WAPL in all settings (Figure S5). Across all chromosomes, 184 genomic bins (100kb-length) containing 217 genes in the unstimulated naïve KO T cells occupied a different compartment when compared to WT T cells (Figure S5). These switch events demonstrated a bias from compartment B to compartment A (70.7% of switch events) (Figure S5).

In the context of lymphopenic stimulation (syngeneic), KO T cells exhibited 303 switch events involving 375 genes when compared to WT T cells. In the allogeneic context, KO T cells demonstrated 413 switch events involving 485 genes when compared to WT T cells (Figure S5). The switch bias was once again in the direction of compartment B to compartment A in syngeneic T cells (60.4% of switch events) and in allogeneic T cells (53.8% of switch events). Additionally, allogeneic T cells had more switch events per chromosome than unstimulated naïve and syngeneic T cells (Figure S5). Interestingly, switched compartment loci tended to congregate towards the ends of chromosomes rather than in the middle or spread evenly throughout. No switch events were observed among contiguous regions of the genome in any of these settings. Overall, 14%, 68%, and 70% of switch events in naive, syngeneic, and allogeneic T cells, respectively, occurred within the previously identified LP-dissimilar regions. While there are coordinated changes in expression and chromatin accessibility genome-wide and most notably at LP-dissimilar regions, chromatin compartmentalization largely remained stable between WT and KO T cells (Table S2).

### WAPL impacts internal structure of TADs and local gene transcription

Stability in global compartment organization throughout the genome does not preclude changes to local genome organization. Chromatin preferentially interacts within locally-distributed and insulated regions called topologically associating domains (TADs) that regulate transcription (Dixon et al., 2012). Thus, we next analyzed the impact of WAPL depletion in TADs. One highly supported mechanism of TAD formation is loop extrusion (Fudenberg et al., 2016) mediated by the ring-shaped cohesin complex and WAPL, which enables TAD dynamics by pro-moting cohesin turnover (Haarhuis et al., 2017). Specifically, genes residing in the same TAD experience coordinated regulation and expression (Dixon et al., 2012; Nora et al., 2012; Shen et al., 2012; Le Dily et al., 2014) while changes to TAD boundaries due to altered CTCF binding influence anomalous gene-enhancer interactions (Flavahan et al., 2016; Lupiáñez et al., 2015). We therefore utilized spectral graph theory to identify the positional boundaries that define TADs (Chen et al., 2016). TADs have been widely suggested to be highly conserved across cell types and conditions in mammalian genomes (Dixon et al., 2012; Nora et al., 2012; Rao et al., 2014), so we pooled our Hi-C data across settings to enable higher resolution binning of the data and reliable detection of TAD boundaries within chromosomes (see Methods).

TAD analysis revealed the emergence of a unique feature in the context of syngeneic and allogeneic T cells in the absence of WAPL - the appearance of ‘corner peaks’. Corner peaks are the enrichment of interaction frequency at domain boundaries seen at the bottom left and top right corners of TADs (Rao et al., 2014). To evaluate differences in corner peak signal between WT and KO T cells, we established a local neighborhood around the corners of each TAD and determined the average number of observed contacts in each neighborhood (Figure 3A). On aggregate, we observed that corner peaks became more pronounced upon WAPL depletion in syngeneic and allogeneic T cells (Figure 3B and Figure S6), consistent with the known activity of WAPL to limit loop extension (Haarhuis and Rowland, 2017; Haarhuis et al., 2017). Further, several studies have demonstrated that corner peaks are associated with a longer residence time of the cohesin complex at TAD borders (Haarhuis et al., 2017; Schwarzer et al., 2017; Szabo et al., 2019). We did not, however, observe this rise in corner peak abundance in the KO unstimulated naïve T cells, but rather a decrease (Figure 3B and Figure S6), suggesting that the development of corner peaks in stimulated T cells might be a consequence of their proliferation following activation.

**Fig. 3:**
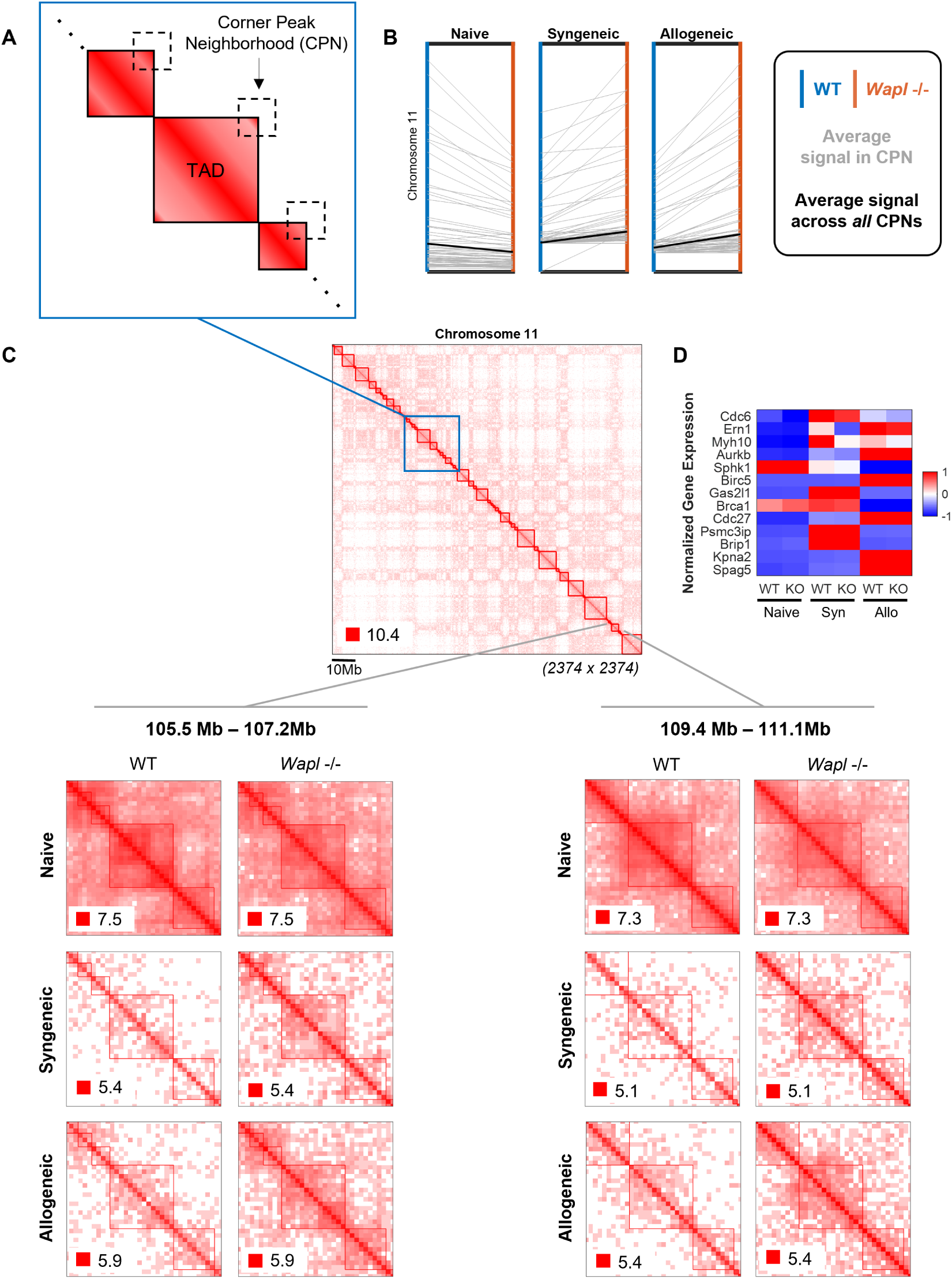
Comparison of internal TAD organization between WT and KO T cells. (A) Schematic of TADs illustrating how ‘corner peaks’ – interactions between opposite TAD boundaries – were characterized in terms of their local neighborhoods. The nonzero mean of absolute read counts in each local corner peak neighborhood (CPN) was used to define the strength of TAD boundary interactions. (B) Change in corner peak signal from WT (blue axis) to KO (orange axis) for each TAD (individual gray lines) on Chromosome 11. The average change in corner peak signal between the two conditions is plotted as a solid black line in each panel to capture the overall trend across settings. TADs and their corner peak signals are represented at 50kb resolution. (C) (Top center) Chromosome 11 observed contact map pooled across samples at 50kb resolution. (Bottom left) TAD region of Chromosome 11 extending from 105.5 Mb to 107.2 Mb capturing corner peak increase from WT to KO most notably in the syngeneic and allogeneic settings; differential expression of gene *Wipi1* occurs in the centermost TAD in this region. (Bottom right) TAD region of Chromosome 11 extending from 109.4 Mb to 111.1 Mb capturing corner peak increase from WT to KO; differential expression of gene *Cdc42ep4* occurs in the centermost TAD in this region. (D) Heatmap of differentially expressed cell cycle genes residing on Chromosome 11.

When compared against a background set of interactions – surrounding interactions occurring at a comparable distance to opposing TAD boundaries - we found that the rise in corner peak signal from WT to KO in syngeneic and allogeneic T cells was statistically significant in a large proportion of TADs (p *<* 0.01, Table S3). We next determined whether there was any functional significance to the observed corner peak dynamics. Syngeneic T cells harbored the most significantly differentially expressed genes (DEGs) (*|*log_2_FC*| ≥* 1, p *≤* 0.01) between WT and KO out of all settings (297 in naïve, 566 in syngeneic, and 46 in allogeneic) and had a particularly high concentration of DEGs on chromosome 11 (58 DEGs, Figure S4). Given this functional dissimilarity, we further investigated intra-TAD interactions on Chromosome 11 at 50kb resolution (Figure 3C) and present an earlier iteration of this analysis on Chromosome 7 at a lower 100kb resolution in Figure S6.

We were particularly interested in identifying regions where differential expression overlapped with increases in corner peak signal. We noted statistically significant corner peak increases in 3 naïve, 47 syngeneic, and 50 allogeneic TADs upon WAPL deletion across Chromosome 11, respectively. Of the TADs exhibiting these corner peak increases, two contained DEGs in naïve T cells, 10 in syngeneic T cells, and three in allogeneic T cells. We highlighted two such regions on Chromosome 11 (Figure 3C, bottom left and right). The first region we observed ranged from position 105.5Mb to 107.2 Mb on Chromosome 11 and encompassed a TAD exhibiting a significant corner peak signal increase most notably in the syngeneic and allogeneic contexts. Furthermore, this TAD contains the gene, *Wipi1*, which exhibited a 2.8-fold increase in expression from WT to KO in syngeneic cells yet a less significant 0.8-fold increase in expression from WT to KO in both naïve and allogeneic T cells. Another stand-out TAD within region 109.4Mb to 111.1Mb on Chromosome 11 demonstrated differential expression of *Cdc42ep4* in syngeneic T cells with a 1.5-fold increase in expression upon WAPL deletion. Overall, Chromosome 11 contained 25 DEGs in naïve T cells, 58 DEGs in syngeneic T cells, and 4 DEGs in allogeneic T cells dispersed across several TADs.

### Impact of WAPL on the Cell Cycle Gene Network

WAPL is critical for sister chromatid cohesion and loop extrusion dynamics. These processes correlate with cellular proliferation and cell cycling, and because we observed notable changes in corner peaks of Chromosomes 7 and 11 only in the context of syngeneic and allogeneic settings in the absence of WAPL, we next explored changes in cell cycle genes and the TADs that these genes reside in. One gene-rich region in Chromosome 7, extending between positions 60Mb and 70Mb on the chromosome, exhibited considerable intra-TAD reorganization in the KO T cells when compared to WT unstimulated (Figure S7D), syngeneic (Figure S7E), and allogeneic (Figure S7F) T cells. This region contains cell cycle genes (Dolatabadi et al., 2017; Inaba et al., 2018): *Fanci*, *Prc1*, and *Blm*. These cell cycle genes (*Fanci*, *Prc1*, and *Blm*) were not significantly differentially expressed, as they did not meet the cutoff of *|*log_2_FC*| ≥* 1. However, there were seven non-cell cycling genes (*Mesp2*, *Arpin*, *Fes*, *Homer2*, *Saxo2*, *Cemip*, and *Arnt2*) directly upstream and downstream of the cell cycle genes in this region that were significantly differentially expressed. These data suggest that in the unstimulated KO T cells, these three cell cycle genes on Chromosome 7 were not differentially expressed despite the changes in intra-TAD interactions when compared to WT T cells.

Similar to the unstimulated context, in the syngeneic T cells, there was a differential expression of non-cell cycle genes upstream and downstream of cell cycle genes in this region (*Agbl1*, *Isg20*, *Can*, *Hapln3*, *Ribp1*, *Mesp2*, *Anpep*, *Fes*, *Slc28a1*, *Homer2*, *Adamtsl3*, and *Tmc3*). In allogeneic T cells, non-cell cycle genes (*Agbl1*, *Can*, *Rhcg*, and *Adamtsl3*) once again were differentially expressed. Thus, in the absence of WAPL, the highlighted region of Chromosome 7 containing the three cell cycle genes did not dynamically change but demonstrated significant changes in the expression (*|*log_2_FC*| ≥* 1, p *≤* 0.05) and internal structural rearrangement of genes in their vicinity. No other cell cycle genes on Chromosome 7 demonstrated differential expression.

Chromosome 11 contained 13 differentially expressed cell cycle genes (Figure 3D). Of these 13, only one gene (*Gas2l1*) was significantly differentially expressed between naïve WT and KO T cells, two genes (*Myh10* and *Sphk1*) between syngeneic WT and KO T cells, and none in the allogeneic setting. We did not however observe an overlap between the differential expression of these genes and TADs with altered internal TAD structure. While WAPL depletion did not disrupt the cell cycling transcriptional program within unstimulated and stimulated contexts much on Chromosome 11, the expression of these genes was quite different between contexts (Figure 3D).

While we did not see a change in the three cell cycle genes noted above in the analysis of Chromosome 7 nor significant coupling of cell cycle differential expression and corner peak dynamics on Chromosome 11, WAPL is known to regulate cell cycling so we next explored the entire breadth of cell cycle genes throughout the genome. To this end, we extracted and stitched together 141 Hi-C genomic bins that corresponded to a curated set of 170 cell cycle genes genome-wide, generating a Hi-C-derived 5C contact matrix (see Methods). 5C, like Hi-C, is a derivative of the original chromosome conformation capture technique (Dekker et al., 2002), and is useful in identifying interactions among select genomic regions that bear relationship to one another (Dostie et al., 2006). In unstimulated naïve T cells, we observed that the connectivity of the cell cycle network decreased in the absence of WAPL, as determined by the element-wise Pearson correlations moving toward zero (Figure 4A). In syngeneic and allogeneic T cells, however, connectivity appeared to strengthen in the absence of WAPL, as demonstrated by the correlation tending towards ± 1 (Figure 4A).

**Fig. 4:**
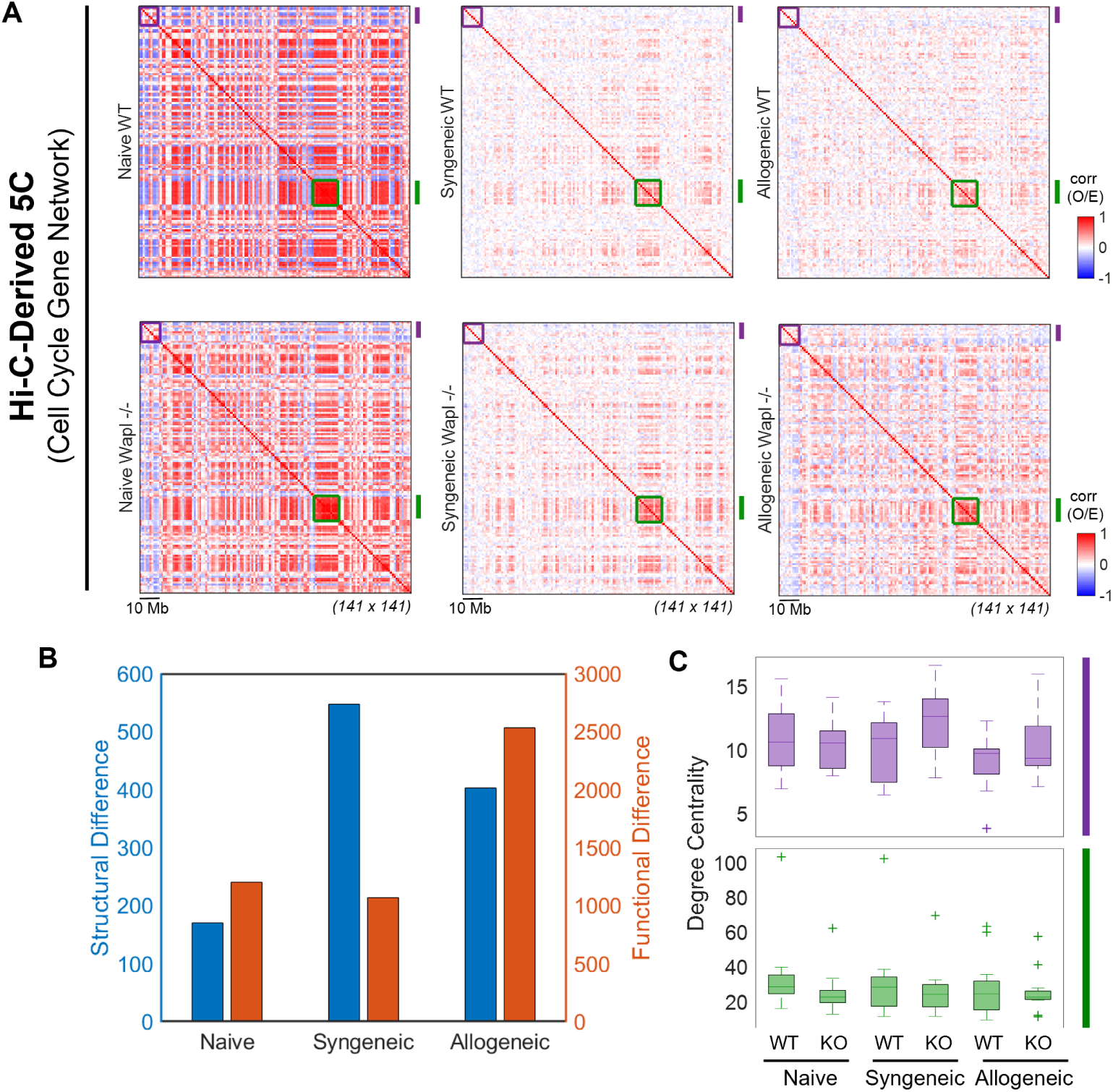
Cell cycle gene network across T cells. (A) Hi-C-derived 5C contact maps representing the cell cycle gene network. Rows and columns correspond to genomic bins across all chromosomes that contain cell cycle genes. Maps are shown at 1-Mb resolution as a Pearson correlation of normalized (observed/expected) contacts. The purple and green boxes highlight subgroups of interest that are assessed in C. The purple subgroup is comprised of 10 loci (1Mb length) ranging non-contiguously from 18 Mb to 197 Mb and containing 10 cell cycle genes. The cell cycle gene loci in WT and KO T cells in each subgroup is comprised of 12 loci (1Mb length) ranging non-contiguously from 1,489 Mb to 1,606 Mb and containing 16 cell cycle genes. (B) Structural and functional differences between all cell cycle gene loci in WT and KO T cells in each setting. Difference is measured as the Frobenius norm of the difference between WT and KO data (see Methods). The left y-axis (blue) reflects the Frobenius norm of the difference between WT and KO Hi-C matrices (measure of structural change) in each setting and the right y-axis (orange) reflects the Frobenius norm of the difference between WT and KO gene expression vectors (measure of functional change) in each setting. (C) Degree centrality (i.e. the row sum of a matrix) of subgroups of interest.

The connectivity (structure) of cell cycle genes changed between the WT and KO T cells in all settings, but the maximal change was noted in the syngeneic setting (Figure 4B). However, changes in the structure of the cell cycle network did not trend with the function (expression). The expression (function) of the cell cycle gene network changed the most between WT and KO T cells in the context of allogeneic setting (Figure 4B). The genome-wide structural analysis of the cell cycle gene network highlighted two subgroups of highly connected genes (shown in purple and green boxes in Figure 4A). One of these subgroups negatively correlated with most of the network (Figure 4A, purple box) while the other demonstrated positive correlation with most of the network (Figure 4A, green box) in all three settings. The measure of interconnectivity within these two subgroups, in terms of degree centrality, is shown in Figure 4C. Ultimately, while gene expression of cell cycle genes *Fanci*, *Prc1*, and *Blm*, did not change in the absence of WAPL, other genes in the cell cycle network did change significantly (Table S4). Five cell cycle genes were up-regulated in the absence of WAPL in unstimulated naive T cells. By contrast, 12 cell cycle genes were down-regulated and 4 up-regulated between WT and KO T cells in the syngeneic context. In the context of allo-antigen stimulation, only one gene was down-regulated.

### WAPL-induced changes in genome structure alter T cell gene expression

We next determined whether the changes in the chromatin architecture related to WAPL deficiency in T cells affected genome function. Three-dimensional genome structural changes have been suggested to affect T cell development (Seitan et al., 2011). Whether *a priori* disruption of the cohesin ring affected T cell function, however, is not clear. The transgenic mice with conditional *Wapl* KO in T cells displayed normal birth and growth rates and generated enough mature naïve T cells. To better analyze the developmental impact of WAPL deficiency on T cell development, we next analyzed the thymii from 8–10-week-old WT and *Wapl* KO littermates. The thymocytes from the *Wapl* KO animals showed significant changes when compared to the WT littermate controls. Specifically, they showed reduction in the total number of thymocytes, double positive (DP), and CD8 SP thymocytes (Figure 5A and Figure S8A).

**Fig. 5:**
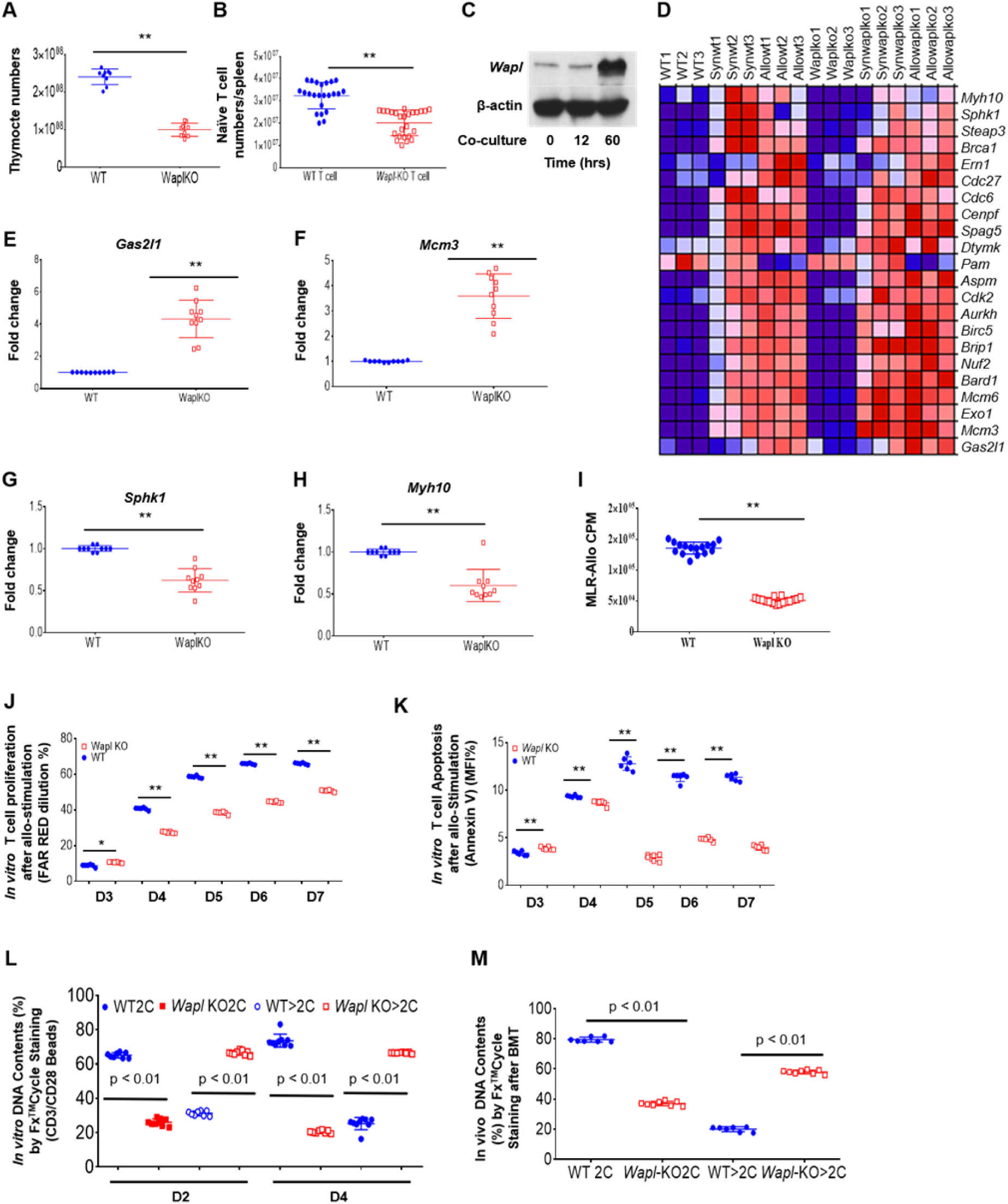
WAPL deficiency impairs T cell development, proliferation, and cell cycling. (A) Total numbers of thymocytes isolated from WT and *Wapl* KO thymii were counted. Data were combined from WT (9) and *Wapl* KO (10) mice (mean ± SEM). (B) Total T cell numbers of spleens from WT and *Wapl* KO mice were counted. Data were combined from WT (23) and *Wapl* KO (26) mice (mean ± SEM). (C) WAPL protein expression was upregulated in T cells when co-cultured with allogeneic DCs in vitro. A representative Western blot was shown. (D) mRNA expression of cell cycle genes was analyzed through GSEA. Gene sets of WT and *Wapl* KO T cells from naïve, syngeneic and allogeneic conditions were sorted by enrichment scores (ESs). All groups were in biological triplicates. (E-H) According to ESs in (D), the mRNA expressions of the top or bottom two scored genes were analyzed by quantitative real-time PCR. Data were combined from three independent experiments (mean ± SEM). (I) H3-TdR incorporation in WT and *Wapl* KO T cells were analyzed by mixed lymphocyte reaction when stimulated by allogeneic DCs for 4 days. Data were combined from three independent experiments (mean ± SEM). (J) Cell proliferation was analyzed by Far-Red dilution for either WT or *Wapl* KO T cells when stimulated with allogeneic DCs for up to 7 days. Data were combined from two independent experiments (mean ± SEM). (K) Cell apoptosis was analyzed by FACS after Annexin V staining for WT and *Wapl* KO T cells when stimulated with allogeneic DCs for up to 7 days. Data were combined from two independent experiments (mean ± SEM). (L) After WT and *Wapl* KO T cells were stimulated with CD3/CD28Ab for 2-4 days *in vitro*, cells were stained with FxCycle™Far-Red, DNA contents were separated as 2C and *>*2C by flow cytometry. Combined data were obtained from three independent experiments (mean ± SEM). (M) WT and *Wapl* KO T cells were isolated from recipient spleens on days 7 after allogeneic BMT *in vivo* and were stained with FxCycle™Far-Red, DNA contents were separated as 2C and *>*2C by flow cytometry. Combined data were obtained from two independent experiments (mean ± SEM).

We next analyzed the secondary lymphoid organ (spleens) for peripheral T cell subsets. *Wapl* KO littermates demonstrated lower numbers of total T cells in the spleen (Figure 5B) and lower numbers of CD3^+^CD4^+^ and CD3^+^CD8^+^ T cells (Figure S8B and C) when compared to WT littermates. During the *β*-selection checkpoint in the thymus, the *β* chain of the T cell receptor rearranged by the thymocytes must retain the structural properties allowing it to be presented on the surface of the thymocytes. We therefore screened for TCR*β*^+^ CD4^+^ and CD8^+^T cells in the spleen and found that the *Wapl* KO mice demonstrated lower numbers of TCR*β*^+^ CD4^+^ and CD8^+^ T cells in the spleen when compared to WT littermates (Figure S8D and E). The splenocytes from the KO animals also demonstrated lower numbers of CD3^+^CD69^+^, CD4^+^CD69^+^, CD8^+^CD69^+^ and CD4^+^CD25^+^ T cells (Figure S8A-D).

Because WAPL deficiency altered chromatin architecture, we next analyzed whether these structural changes were associated with changes in gene expression and proliferation, at baseline and following *in vivo* stimulation. To this end, we first determined whether *Wapl* expression itself changed in T cells following stimulation. Consistent with our previous observation, we observed significant upregulation of WAPL protein in T cells upon co-culture with allogeneic DCs for 60 hours (Figure 5C) (Sun et al., 2019, 2015). To determine the impact of this structurefunction relationship, we performed gene set enrichment analysis (GSEA) for the cell cycle gene network highlighted in our earlier Hi-C-derived 5C analyses (Figure 4). We found that cell cycle genes were differentially regulated between WT and WAPL-deficient T cells (Figure 5D). We verified results with real-time quantitative PCR. *Gas2l1*, a gene induced upon growth arrest (Goriounov et al., 2003), and *Mcm3* that increases as cells progress from G0 into the G1/S phase and regulates the cell cycle (Tsuruga et al., 1997), were significantly upregulated in WAPL-deficient T cells compared to WT T cells (Figure 5E and F). In contrast, *Sphk1*, a gene that plays a key role in TNF-a signaling and the canonical NF-*κ*B activation pathway important in inflammatory, apoptotic, and immune processes (Alvarez et al., 2010), and *Myh10*, a gene required for completion of cell division during cytokinesis (Kim et al., 2005; Straight et al., 2003) were both downregulated in WAPL-deficient T cells (Figure 5G and H). These observations demonstrated that changes in genome structure related to WAPL deficiency in T cells promote differential gene expression that suppresses apoptosis, cell proliferation and cell cycle progress.

To further investigate the impact of WAPL deficiency on T cell cellular responses, we stimulated WAPL-deficient and WT T cells *in vitro*, either with allo-antigen stimulation in mixed lymphocytes reaction (MLR) by co-culturing T cells with allogeneic DCs for four days or with CD3/CD28Ab for two or three days and pulsed with H3-TdR. The WT T cells demonstrated significantly greater H3-TdR incorporation when compared with *Wapl* KO T cells suggesting that WAPL deficiency caused reduced proliferation (Figure 5I and Figure S9E). T cell proliferation was directly assessed *in vitro* using a dye dilution assay. Specifically, we utilized Cell-Trace™FarRed dilution to avoid the interference with GFP fluorescence in *Wapl* KO T cells. The *Wapl* KO T cells demonstrated reduced dye dilutions when compared to WT T cells upon stimulation by either allogeneic DCs or CD3/CD28ab (Figure 5J and Figure S8F). It is possible that cell death from apoptosis could contribute to the reduction in T cell expansion related to WAPL deficiency. We therefore also determined apoptosis after stimulation of both WT and *Wapl* KO T cells. The WAPL-deficient T cells also showed significantly decreased apoptosis when compared to littermate WT T cells (Figure 5K). These data demonstrate that the absence of WAPL altered T cell proliferation and apoptosis following *in vitro* stimulation.

The interaction between cohesin and WAPL plays an essential role in maintaining chromosome structure and separation of sister chromatids during mitosis and cell proliferation (Peters and Nishiyama, 2012). With the coordinated changes in expression of cell cycling genes and genomic architecture in the absence of WAPL, and the reduction in proliferation, we next examined cellular mechanisms underlying the reduced proliferative capacity of WAPL-deficient T cells following *in vitro* and *in vivo* stimulation. Specifically, we explored the hypothesis that altered gene expression of cell cycling genes from the change in genomic architecture related to WAPL deficiency leads to a reduction in cell cycling. To assess kinetic cell cycling we utilized flow cytometry analyses of DNA content with FxCycle™Far Red Stain to avoid interference from the GFP fluorescence in *Wapl* KO T cells. Purified WT or *Wapl* KO T cells were stimulated with CD3/CD28T cell activator dynabeads and analyzed for 2C and *>*2C populations. *Wapl* KO T cells demonstrated significantly lower 2C percentages but higher *>*2C percentages when compared with WT T cells at several time points after CD3/CD28 stimulation (Figure 5L and Figure S10A). We next analyzed whether the effect on cell cycling was observed *in vivo*. To this end, we once again utilized the allogeneic bone marrow transplantation (BMT) model system, where the host splenocytes were harvested and analyzed for WT or *Wapl* KO donor T cells as above. As shown in Figure 5M and Figure S9B, the *in vivo* stimulation of WT and *Wapl* KO T cells demonstrated similar differences as from *in vitro* stimulation. These data indicated that *Wapl* KO T cells had cell cycle deficiency, which impairs T cell proliferation and alters T cell function *in vitro* and *in vivo*, consistent with cell cycle gene analyses by Hi-C derived 5C (Figure 5), GSEA, and qPCR analyses (Figure 5D-H).

### WAPL deficiency in T cells improves survival after allogeneic BMT

Mature T cells in the allogeneic donor T cells are the principal mediators of GVHD, a major cause of mortality after allogeneic BMT. Because WAPL regulated mature T cell responses following *in vitro* and *in vivo* allo-antigen stimulation, we next determined whether this has a clinical impact on GVHD severity following experimental allo-BMT. To this end, we once again utilized the MHC mismatched B6 (H2) → BALB/c (H2d) mouse model where the congenic B6 animals served as the syngeneic controls. BALB/c recipient mice were lethally irradiated (800 cGy total body irradiation, split dose) and transplanted with T cell-depleted WT BM from B6 donors along with purified mature T cells from the spleen of either WT or *Wapl* KO B6 animals(Sun et al., 2015). The recipient animals were monitored for survival and GVHD severity as described in Methods.

We first determined whether WAPL expression changed after allo-BMT. Consistent with the *in vitro* allo-antigen stimulation shown in Figure 5C, WT cells demonstrated higher expression of WAPL protein in donor T cells harvested from recipient spleens 7 days after allogeneic BMT when compared to syngeneic controls (Figure 6A). Survival analysis demonstrated that all the syngeneic animals survived, but the allogeneic animals that received WT T cells died with signs of severe clinical GVHD (Figure 6B). In contrast, allogeneic animals that received WAPL-deficient T cells showed significantly improved survival (53% versus 17%; p *<* 0.01) (Figure 6B) and reduced clinical severity of GVHD (p *<* 0.01) (Figure 6C). We confirmed the reduction in GVHD with detailed histopathological analyses of GVHD target organs including the liver, gastrointestinal (GI) tract, and skin. As shown in Figure 6D, allogeneic animals that received T cells from WAPL-deficient donors had significantly reduced histopathological GVHD in the liver (p *<* 0.01), GI tract (SI and LI) (p *<* 0.05) and skin (p *<* 0.05) on day +21 after BMT. Consistent with reduced mortality, the recipients of allogeneic *Wapl* KO T cells demonstrated reduced serum levels of proinflammatory cytokines such as IFN-*γ* and TNF-*α* when compared with WT T cell recipients (Figure 6E and F).

**Fig. 6:**
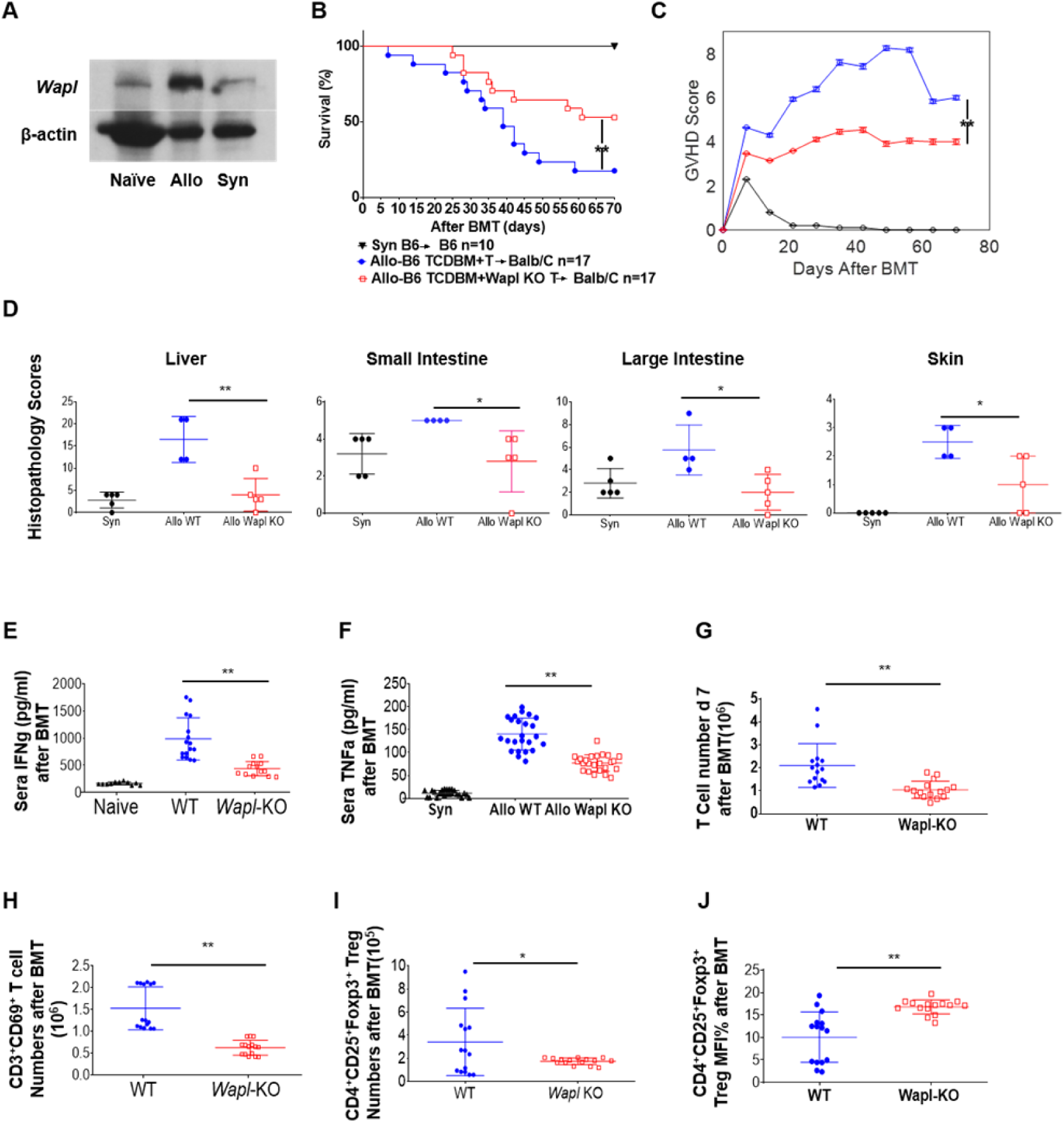
WAPL-deficient T cells mitigate GVHD severity in allogeneic BMT. (A) *Wapl* expression was upregulated in T cells on day 7 after allogeneic BMT. A representative Western blot was shown. (B) Allogeneic MHC mismatched B6 (H2b) into BALB/c(H2d) GVHD model. Survival percentages were recorded daily. P-values were obtained using the log-rank test based on comparisons between animals that received either TCD BM plus WT T cells (blue circles) or TCD BM plus *Wapl* KO T cells (red squares). Combined data were from 2 independent experiments. (C) GVHD score was assessed by a standard scoring system as before (Sun et al., 2015). The Mann-Whitney U test was used for the statistical analysis of clinical scores between animals that received either TCD BM plus WT T cells (blue circles) or TCD BM plus *Wapl* KO T cells (red squares). Combined data were from 2 independent experiments. (D) Histopathological analysis of the B6 into BALB/c BMT model. Bowel (small and large intestine), liver, and skin were obtained after BMT on day 21. GVHD scores were from 2 independent experiments (mean ± SEM). (E and F) Sera were collected from recipient mice on day 21 after allogeneic BMT as in (B), the concentrations of IFN-*γ* and TNF-*α* were measured by ELISA. Significant lower concentrations of INF-*γ* and TNF-*α* were detected in the sera collected from mice transferred with *Wapl* KO T cells. Combined data were from two independent experiments (mean ± SEM). (G) *In vivo* T cell expansion was determined by isolating transferred WT and *Wapl* KO in BALB/c recipients on day 7 after allogeneic BMT. Data were combined from three independent experiments (mean ± SEM). (H) Transferred donor T cells (WT and *Wapl* KO) were isolated from recipient’s spleens on day 7 after allogeneic BMT and stained for CD3^+^ and CD69^+^. The absolute numbers of activated *Wapl* KO T cells (CD3^+^CD69^+^) were significantly lower than WT T cells. Data were combined from three independent experiments (mean ± SEM). (I and J) Donor T cells were isolated as in (H) on day 7 after allogeneic BMT and stained for CD4^+^ 25^+^ FoxP3^+^ for identifying T regulatory cells. The absolute numbers of *Wapl* KO Treg cells were significantly lower than WT Treg cells (I), although the percentages were the same between WT and *Wapl* KO T cells (J). Data were combined from three independent experiments (mean ± SEM). P-values were obtained using unpaired t-test.

The allogeneic animals that received WT or KO T cells demonstrated *>*98% donor engraftment on day 21, ruling out mixed chimerism or engraftment as a cause for reduction in GVHD. Furthermore, consistent with above results, *Wapl* KO T cells showed significantly reduced expansion when compared with WT donor T cells (Figure 6G) in the recipient spleens harvested 7 days after BMT, suggesting that the reduction in GVHD was a consequence of reduced expansion of allo-antigen stimulated T cells. Consistent with this, the allogeneic *Wapl* KO T cells demonstrated fewer absolute numbers of CD3^+^CD69^+^ cells (Figure 6H) and regulatory T cells (despite a higher percent) when compared to WT T cells (Figure 6I and J).

## Discussion

The cohesin complex and its regulators, such as WAPL, are critical determinants of 3D genome architecture, which regulates replication, repair, and transcriptional processes (Cuadrado and Losada, 2020; Misteli, 2020; Nasmyth and Haering, 2009; Remeseiro et al., 2013). Given the paucity of data on the genomic organization of mature T cells and its impact on T cell function *in vivo*, we describe the genome architecture of mature T cells following *in vivo* lymphopenic and allogeneic stimulation.

Prior to the availability of Hi-C methods, structural features of the T cell genome following *in vitro* stimulation were explored in a seminal paper by Kim *et al*. (Kim et al., 2004). Since then, other studies have built on it with the advent of Hi-C techniques following *in vitro* stimulation of T cells with anti-CD3/CD28 (Yang et al., 2020; Burren et al., 2017; Bediaga et al., 2021).

We now expand on those studies by exploring the changes following *in vivo* homeostatic and antigen-driven stimulation. Previously, chromosome 6 in differentiating T cells was explored in its entirety, though global spatial characterization was limited to chromosome territories (Kim et al., 2004). Here, we define the entire genome architecture of mature T cells following *in vivo* activation, including local structures like TADs. We further connect those changes to genomewide functional changes in gene expression and T cell responses. Hu *et al*. demonstrated a key role for BCL11B in the development of T cells and associated genome structure changes (Hu et al., 2018), but did not characterize the relevance of structural integrity to mature T cell functions. Our data focus on the genomic landscape of mature T cells. We define a mechanistic role for genomic architecture in regulation of gene expression, cellular function, and biological responses *in vivo* that impact clinically relevant disease states such as GVHD. We determine that differences in 3D genome architecture are a consequence of the cellular state of activation, and that its disruption, by altering the function of the cohesin complex, regulates mature T cell functions in response to allogeneic-stimulation.

We explored the mechanistic relevance of genome structure to function (gene expression) and to the cellular functions and biological impact of mature T cells by deleting WAPL, a key regulator of genome structure. WAPL deletion led to reduction of long-range interactions in the baseline unstimulated state of naïve T cells. However, following lymphopenic (syngeneic) and antigen (allogeneic) activation, there was a greater amplification of longer-range interactions following WAPL deletion. This is consistent with a previous report on WAPL-deficient cell lines (Haarhuis et al., 2017; Liu et al., 2021) and with the notion of extruded DNA loops beyond CTCF barriers (Allahyar et al., 2018). These data suggest that WAPL alters genomic architecture, at baseline and after activation of T cells.

The *in vivo* role for WAPL-mediated regulation of chromatin structure is crucial for embryonal development (Tedeschi et al., 2013). We now extend these studies and demonstrate that WAPL also regulates *in vivo* immune responses mediated by T cells. T cell-specific WAPL-deficient animals developed normally despite the T cells showing genomic structural changes at homeostasis. Upon stimulation, WAPL-deficient T cells demonstrated mitosis defects and more axial structures during interphase suggesting that they exit mitosis with less intact cohesin. However, it is important to note that the absence of WAPL neither caused a complete loss of development of T cells nor a total shutdown of mature T cell proliferation. The T cells developed and proliferated in the absence of WAPL, albeit at a much lower level, the reasons for which remain unclear. One possible explanation is that separation of chromatids during mitosis in T cells may be independent, or only partially dependent, on WAPL, based on their strength and duration of stimulation/activation (Oliveira and Nasmyth, 2013; Srinivasan et al., 2019; Yuen and Gerton, 2018). This notion is consistent with the observation that several cohesinopathies in humans are caused by mutations in various components of cohesin complex and yet do not appear to cause T cell defects (Piché et al., 2019; Remeseiro et al., 2013; Singh and Gerton, 2015). Future studies may determine the role of WAPL and disruption of cohesin in thymopoiesis and in regulation of T cells in secondary lymphoid organs.

The structural changes in our study reflect a polyclonal response from a combination of alloreactive/lymphopenia induced proliferating cells, and the non-proliferating mature T cells. The T cells therefore might be in various stages of early/mid G1, S, G2, M phases. Thus, the genome contact frequency and associated structural changes are reflective of the T cells in these various stages after stimulation. T cells from naïve, syngeneic, and allogeneic settings demonstrated significant changes in internal TAD structure, which were consistent despite the polyclonal nature of the T cell subsets. Nonetheless, while the genome architectures were significantly different, whether these are the direct cause or a consequence of proliferation differences cannot be definitively ascertained. Furthermore, whether antigen-specific T cell responses vary based on the type of antigen cannot be determined from our study. However, our assessment of polyclonal responses is akin to biologically and clinically relevant situations such as allogeneic transplantation.

Our data collectively demonstrate for the first time, to our knowledge, that altering genomic structure *a priori*, at baseline, regulates T cell gene and cellular functions. Mechanistically, the data show that WAPL alters chromatin architecture at cell cycling genes and thus links genome structure and function. While we only validate select cell cycling genes concomitant with changes in genome structure in this study, it is possible that other cell cycling genes might be involved in the proliferative defects we observe and future studies will need to validate these in a systematic manner. Future studies may also benefit from integrating single-cell Hi-C and single-cell RNA-seq to refine observations, as features at the TAD and sub-TAD levels observed in this study via bulk Hi-C are a population average and could be variable and distinct between various T cell subsets that develop and differentiate after stimulation. Regardless, our data demonstrate that disruption of 3D chromatin architecture by WAPL can directly regulate gene expression and cellular function of T cells in a physiologically and clinically relevant disease context.

Importantly, we introduce a simple yet robust approach for evaluating TADs and their internal structure. Due to the experimental limitations of performing Hi-C in an *in vivo* setting we obtained a lower number of cells and thus mappable reads than what Hi-C performed on cell lines or an *in vitro* setting would typically yield. Consequently, to reliably investigate TAD and sub-TAD level features, we pooled data across all our samples allowing us to bin our data at a higher resolution. From this pooled matrix, we were able to determine TAD boundaries and subsequently superimpose them onto each sample’s raw Hi-C contact map. Informed by this common TAD backbone, we were able to perform targeted analysis of the internal TAD structure in each sample and evaluate how sensitive the backbone and its internal organization is to perturbations across the different settings.

There are no reported cases of isolated germline WAPL mutations in humans (Gard et al., 2009; Piché et al., 2019; Remeseiro et al., 2013; Singh and Gerton, 2015). This is likely because of its critical role during embryonic development. However, somatic mutations in WAPL have been linked to epithelial carcinogenesis (Waldman, 2020). Our study demonstrates that WAPL deficiency in T cells did not cause a profound defect in development of T cells, nor cause T cell malignancy. Thus, WAPL may play a nuanced role in different cell subtypes, depending on their developmental stage, context, and stimulation. Future studies will need to carefully assess the biological implications of WAPL deficiency on T cell subsets and other immune cells. Because WAPL-deficient T cells demonstrated a reduction in GVHD, it is tempting to speculate whether targeting it uniquely in T cells might be a novel strategy to mitigate immunopathologies such as GVHD, allograft rejection, or autoimmunity. Such a strategy may be feasible with emerging gene editing strategies for adoptive transfer of T cells, however, the viability of the strategy to delete WAPL clinically will need significant further investigation. At a broader level, our data provide a proof of concept for the notion that targeting 3D genomic architecture may be a therapeutic strategy that can be potentially harnessed for directly modulating *in vivo* disease processes.

## Author Contributions

Conceptualization, P.R. and I.R.; Methodology, Y.S., G.A.D. W.M.,I.R., P.R.; Software, G.A.D., S.R.; Validation, Y.S., G.A.D, I.R., P.R. ; Formal Analysis, Y.S., G.A.D; Investigation, Y.S., G.A.D, L.A.M., S.R., K.O.W., D.P. K.S., W.M.; Data Curation, G.A.D.; Writing – Original Draft, Y.S., G.A.D., I.R., P.R.; Writing - Review Editing, Y.S., G.A.D., L.A.M., S.R., K.O.W., D.P. K.S., L.L., W.M., I.R., P.R.; Visualization, Y.S., G.A.D., I.R., P.R.; Supervision, I.R., P.R.; Project Administration, I.R., P.R.; Funding Acquisition, I.R., P.R.

## Acknowledgements

This work is supported by the University of Michigan Genome Science Training Program (GSTP) Fellowship funded by NHGRI under Award Number 5T32HG000040-27 to G.A.D. as well as awards R01HL152605, P01HL149633, and R01CA217156 to P.R.

## Resource Availability

### Lead contact

Further information and requests for resources, reagents and strains should be directed to and will be fulfilled by the lead contact, Pavan Reddy or Indika Rajapakse.

### Materials availability

Mouse strain generated in this study is available from the lead contact with a completed Materials Transfer Agreement.

### Data and code availability

RNA-seq and Hi-C data have been deposited at GEO and BioProject, respectively, and are publicly available. Accession numbers are listed in the key resources table. This paper does not report original code. Any additional information required to reanalyze the data reported in this paper is available from the lead contact upon request.

## Experimental Model and Subject Details

### Mice

B6 (H2b, CD4.1 and CD45.2), BALB/c (H2d, CD45.1 and CD45.2), Rosa26-floxed STOP-Cas9 knockin (B6J) and CD4CRE (B6) mice were purchased from The Jackson Laboratory and the National Cancer Institute. The ages of the mice used for experiments ranged between 8 and 12 weeks. Mice were housed in sterilized microisolator cages and received filtered water and normal chow.

### Generating conditional *Wapl* KO T cells

We designed 2 sgRNAs targeting exon 2 of the *Wapl* locus which are localized in mouse Chromosome 14 qB (Figure S3) (Ran et al., 2015). Three lines of mice with B6 background were used to generate conditional *Wapl* KO mice in T cells. The first line was generated to carry CRISPR-sgRNAs targeting insert to identify exon 2 in *Wapl* (Figure 1A). The second and third lines are Rosa26-floxed STOP-Cas9-GFP knockin on B6J (The Jackson Laboratory, Stock No:026175) and CD4-CRE transgenic mice (The Jackson Laboratory, Stock No: 017336). Tail DNA was screened for a positive sgRNAs-*Wapl* insert, positive CRE, and a stop signal in loxP-SV40pA x3-loxP cleavage or deletion. The potential *Wapl* KO T cells were sorted for positive GFP (Figure 1B). To confirm the success of conditional in KO in T cells, T cells were processed for SDS-PAGE and detected with anti-*Wapl* antibody. Additionally, total RNAs were isolated from *Wapl* KO and WT T cells and processed for RNA-seq.

### DC isolation and purification

Dendritic cells (DC) were isolated from splenocytes either from WT B6 or BALB/c mice. Single-cell suspensions were prepared according to the manufacturer’s instruction then subjected to CD11c microbead (MACS) staining and positive selection using LS column (Miltenyi Biotec). The purity of enriched CD11c^+^ DC preparation was 85.6-90%.

### T cell isolation and purification, and mixed lymphocyte reaction (MLR)

WT and *Wapl* KO B6 T cells were isolated by negative selection (*>*95% purity) (Pan T Cell Isolation Kit II; Miltenyi Biotec). T cells were co-cultured with B6 WT or BALB/c DCs at a ratio of 40:1 (T cells versus DCs 3 × 105:7.5 × 103) for 96 hours using 96-well flat-bottomed plates (Falcon Labware), or stimulated with Dynabeads T cell activator CD3/CD28 (25 ul/106/ml) for 2 or 3 days respectively. Proliferation was determined by incubating the cells with H3-thymidine (1 Ci/well [0.037 MBq]) for the last 20 or 6 hours, respectively. H3-thymidine incorporation in T cells was counted on a 1205 Betaplate reader (Wallac, Turku, Finland).

### BMT and systemic analyses of GVHD

Bone marrow transplantations (BMTs) were performed as described previously (Sun et al., 2015). The donor T cells (WT B6 or WAPL deficiency) were isolated from spleens and purified by negative selection (using the Pan T cell Isolation Kit II; Miltenyi Biotec). Bone marrow cells from tibia and fibula were harvested and TCD (T cell deletion) BM cells were isolated with positive deletion using anti-CD90.2 microbeads and LS column (Miltenyi Biotec). The recipient BALB/c mice received an 800-cGy total body irradiation on day -1 (split dose) and T cells (1 × 106, either B6 WT or WAPL deficiency T cells, and TCD BM cells (5 × 106, from WT B6 mice) were injected intravenously into the recipients on day 0. The syngeneic B6 control mice received a 1000-cGy total body irradiation on day -1. T cells (2 × 106, isolated from B6 WT mice) and TCD BM cells (5 × 106 from B6 WT mice) were injected intravenously into the recipients on day 0. Mice were housed in sterilized microisolator cages and received normal chow and autoclaved hyperchlorinated drinking water for the first 3 weeks after BMT.

### Immunoblotting

T cell lysates were extracted as previously described (Sun et al., 2015), and 50 to 100 g of protein extract was separated in SDS-PAGE and transferred onto a PVDF membrane (GE Healthcare). The membrane was blocked with 5% nonfat milk for 30 minutes and then incubated overnight at 4°C with the following Abs in 5% nonfat milk: anti-Wapl rabbit polyclonal Ab (1:500 in nonfat milk, Proteintech Cat 16370-1-AP), anti–*β*-actin mouse mAb (1:1,000 in 5% nonfat milk; Abcam, catalog ab8226). After washing 3 times with TBST for 5 minutes, the blot was incubated with specific HRP-labeled secondary Ab, washed again with TBST, and signals generated and visualized using the Enhanced Chemiluminescence Kit (Thermo Scientific, Cat 32106).

### Study approval

Study approval. All animal studies were reviewed and approved by the University Committee on Use and Care of Animals of the University of Michigan, based on University Laboratory Animal Medicine guidelines.

## Method Details

### GVHD and pathology scoring

Survival was monitored daily. The degree of systemic GVHD was assessed by a standard scoring system with four criteria scores: percentage of weight change, posture, activity, fur texture, and skin integrity, and subsequently graded from 0 to 2 for each criterion (maximum index = 10) (Sun et al., 2015). Acute GVHD was also assessed by histopathologic analysis of the ileum and the ascending colon, liver, and ear skin. Specimens were harvested from animals on day 21 after BMT, then processed and stained with hematoxylin and eosin. Coded slides were examined systematically in a blind manner by using a semi-quantitative scoring system to assess the following abnormalities known to be associated with GVHD, small intestine: villous blunting, crypt regeneration, loss of enterocyte brush border, luminal sloughing of cellular debris, crypt cell apoptosis, outright crypt destruction, and lamina propria lymphocytic infiltrate; colon: crypt regeneration, surface colonocytes, colonocyte vacuolization, surface colonocyte attenuation, crypt cell apoptosis, outright crypt destruction, and lamina propria lymphocytic infiltrate; liver: portal tract expansion, neutrophil infiltrate, mononuclear cell infiltrate, nuclear pleomorphism, intraluminal epithelial cells, endothelialitis, hepatocellular damage, acidophilic bodies, mitotic figures, neutrophil accumulations, macrophage aggregates, macrocytosis; skin: apoptosis in epidermal basal layer or lower malpighian layer or outer root sheath of hair follicle or acrosyringium, lichenoid inflammation, vacuolar change, lymphocytic satellitosis. The scoring system denoted 0 as normal, 0.5 as focal and rare, 1.0 as focal and mild, 2.0 as diffuse and mild, 3.0 as diffuse and moderate, and 4.0 as diffuse and severe. Scores were summed together to provide a total score for each specimen (Sun et al., 2015).

### RNA-seq library generation and data processing

Naïve WT and *Wapl* KO T cells, or WT and *Wapl* KO T cells isolated from syngeneic or allogeneic BMT on day 7 were first purified as described previously (Sun et al., 2015). All dead cells were excluded by sorting for far-red fluorescent reactive dye (Invitrogen, Cat. L10120). Then, WT T cells were sorted for PE-CD3 and APC-CD45.2 while *Wapl* KO T cells were sorted for APC-CD3, PE-CD45.2 and positive GFP. Each sample contained pooled T cells from 3-4 mice, and samples in each group were biologically triplicated. RNA was isolated using DNA/RNA mini Kit (Qiangen Cat 80204) by following the RNA isolation procedures. Sequencing was performed by the University of Michigan (UM) DNA Sequencing Core, using the Illumina Hi-Seq 4000 platform, paired-end, 50 cycles and Ribosomal Reduction library prep.

We obtained read files from the UM Sequencing Core and concatenated those into a single FASTQ file for each sample. We checked the quality of the raw read data for each sample using FastQC (version 0.11.3) to identify features of the data that may indicate quality problems (e.g., low quality scores, overrepresented sequences, inappropriate GC content). We used the Tuxedo Suite software package for alignment, differential expression analysis, and post-analysis diagnostics. Briefly, we aligned reads to the reference genome (GRCm38) using TopHat (version 2.0.13) and Bowtie2 (version 2.2.1). We used default parameter settings for alignment, with the exception of: “–b2-very-sensitive” telling the software to spend extra time searching for valid alignments. We used FastQC for a second round of quality control (post-alignment), to ensure that only high-quality data would be input to expression quantitation and differential expression analysis. We used Cufflinks/CuffDiff (version 2.2.1) for expression quantitation, normalization, and differential expression analysis, using GRCm38.fa as the reference genome sequence. For this analysis, we used parameter settings: “–multi-read-correct” to adjust expression calculations for reads that map in more than one locus, as well as “–compatible-hits-norm” and “–upper-quartile–norm” for normalization of expression values. We generated diagnostic plots using the CummeRbund R package. We used locally developed scripts to format and annotate the differential expression data output from CuffDiff. Briefly, we identified genes and transcripts as being differentially expressed based on FDR 0.05, and fold change ± 1.5. We annotated genes with NCBI Entrez GeneIDs and text descriptions. RNA-seq reads in bam format were mapped to the most recent mouse genome (mm10) using the Integrative Genomics Viewer (IGV). The RNA-seq data reported here can be found in the Gene Expression Omnibus (GEO) database with the series accession ID GSE134975.

### Generation of Hi-C libraries for sequencing

The *in situ* Hi-C protocols from Rao *et al*. (Rao et al., 2014) were adapted with slight modifications. For each Hi-C library, approximately 1 x 106 cells were incubated in 250 µl of ice-cold Hi-C lysis buffer (10mM Tris-HCl pH8.0, 10mM NaCl, 0.2% Igepal CA630) with 50µl of protease inhibitors (Sigma) on ice for 30 minutes and washed with 250 µl lysis buffer. The nuclei were pelleted by centrifugation at 2500xg for 5 minutes at 4°C, re-suspended in 50 µl of 0.5% sodium dodecyl sulfate (SDS) and incubated at 62°C for 10 minutes. Afterwards 145 µl of water and 25 µl of 10% Triton X-100 (Sigma) were added and incubated at 37°C for 15 minutes.

Chromatin was digested with 200 units of restriction enzyme MboI (NEB) overnight at 37°C with rotation. Chromatin end overhangs were filled in and marked with biotin-14-dATP (Thermo Fisher Scientific) by adding the following components to the reaction: 37.5 µl of 0.4mM biotin-14-dATP (Life Technologies), 1.5 µl of 10mM dCTP, 1.5 µl of 10mM dGTP, 1.5 µl of 10mM dTTP, and 8 µl of 5U/µl DNA Polymerase I, Large (Klenow) Fragment (NEB). The marked chromatin ends were ligated by adding 900 µl of ligation master mix consisting of 663 µl of water, 120 µl of 10X NEB T4 DNA ligase buffer (NEB), 100 µl of 10% Triton X-100, 12 µl of 10mg/ml BSA, 5 µl of 400 U/µl T4 DNA Ligase (NEB), and incubated at room temperature for 4 hours.

Chromatin de-crosslinking was performed by adding 50 µl of 20mg/ml proteinase K (NEB) and 120 µl of 10% SDS and incubated at 55°C for 30 minutes, adding 130 µl of 5M sodium chloride and incubate at 68°C overnight. DNA was precipitated with ethanol, washed with 70% ethanol, and dissolved in 105 µl of 10 mM Tris-HCl, pH 8. DNA was sheared on a Covaris S2 sonicator. Biotinylated DNA fragments were pulled with the MyOne Streptavidin C1 beads (Life Technologies). To repair the ends of sheared DNA and remove biotin from unligated ends, DNA-bound beads were re-suspended in 100 µl of mix containing 82 µl of 1X NEB T4 DNA ligase buffer with 10mM ATP (NEB), 10 µl of 10 (2.5mM each) 25mM dNTP mix, 5 µl of 10U/µl NEB T4 PNK (NEB), 4 µl of 3U/µl NEB T4 DNA polymerase (NEB), and 1 µl of 5U/µl NEB DNA polymerase I, Large (Klenow) Fragment (NEB).

After end-repair, dATP attachment was carried out in 100 µl of reaction solution, consisting of 90 µl of 1X NEBuffer 2, 5 µl of 10mM dATP, and 5 µl of 5U/µl NEB Klenow exo minus (NEB). The reaction was incubated at 37°C for 30 minutes. The beads were then cleaned for Illumina sequencing adaptor ligation which was done in a mix containing 50 µl of 1X T4 ligase buffer, 3 µl T4 DNA ligases (NEB), and 2 µl of a 15 µM Illumina indexed adapter at room temperature for 1 hour. DNA was dissociated from the bead by heating at 98°C for 10 min, separated on a magnet, and transferred to a clean tube.

Final amplification of the library was carried out in multiple PCRs using Illumina PCR primers. The reactions were performed in 25 µl scale consisting of 25 ng of DNA, 2 µl of 2.5mM dNTPs, 0.35 µl of 10 µM each primer, 2.5 µl of 10X PfuUltra buffer, PfuUltra II Fusion DNA polymerase (Agilent). The PCR cycle conditions were set to 98°C for 30 seconds as the denaturing step, followed by 14 cycles of 98°C 10 seconds, 65°C for 30 seconds, 72°C for 30 seconds, then with an extension step at 72°C for 7 minutes.

After PCR amplification, the products from the same library were pooled and fragments ranging in size from 300-500 bp were selected with AMPure XP beads (Beckman Coulter). The size-selected libraries were sequenced to produce paired-end Hi-C reads on the Illumina HiSeq 2500 platform with the V4 of 125 cycles.

### Hi-C data processing

We generated the Hi-C matrices using Juicer (Durand et al., 2016). Juicer uses BWA-mem to align each paired-end read separately. It then determines which reads can be mapped uniquely and keeps unambiguously mapped read pairs. Each read is assigned to a ”fragment”, determined by the restriction enzyme cut sites and paired reads that map to the same fragment are removed. The Juicer pipeline creates a binary data file (namely, the “.hic” file), which contains Hi-C contacts. The .hic file is imported into MATLAB-compatible variables via our in-house MATLAB toolbox, 4DNvestigator (Lindsly et al., 2021). Centromeric and telomeric regions were removed in the process. Then, Hi-C data were subsequently normalized and binned at 50kb and 100kb resolution. ICE (Imakaev et al., 2012) and “observed over expected” (O/E) normalized contact maps at 100kb resolution were used for chromosome-level analysis of A/B compartments. Observed contact maps at 50kb resolution were used for detection and analysis of TADs. Hi-C data for this study are available through the NCBI BioProject database (accession number: PRJNA608895).

### Integration of Hi-C and RNA-seq data

To integrate our Hi-C and RNA-seq datasets, we first binned measurements at the same resolution. Accordingly, each element in the binned RNA-seq expression vector corresponds to a row (or column) of the Hi-C contact matrix that captures the same set of genomic loci. Genomic bins spanning a given region contain measurements reflecting the sum of its parts, where expression values for genes sharing the same bin are summed and contact frequencies for loci occupying the same bin are summed. This allows us to directly compare the transcriptional and conformational behavior across regions of the genome.

### Frobenius Norm

The Frobenius norm describes the size or volume of a matrix (Strang et al., 1993). As such, the Frobenius norm can be used as a measure of variance or variability in data (Drineas and Mahoney, 2018). For matrix A of dimension m, that is A *∈*R^m^*^×^*^n^, the mathematical definition of the Frobenius norm is as follows:

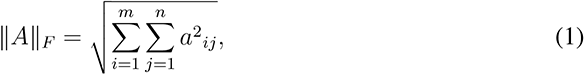

where *a_ij_* is an element of the data matrix *A*. Equivalently this expression can be written as,

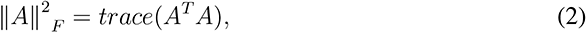

where *trace* is the sum of the diagonal elements of a matrix (Eldén, 2007). When comparing two sets of data, the notation can is modified as follows:

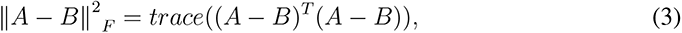

where *A* and *B* are matrices of the same dimensions. The Frobenius norm of a vector is equivalent to the Euclidean norm.

### Larntz-Perlman Procedure

The Larntz-Perlman (LP) procedure is a method designed to test the equivalence of correlation matrices (Larntz and Perlman, 1985). We applied the LP procedure as described in the 4DNvestigator Toolbox (Lindsly et al., 2021) to compare Hi-C contact matrices between WT and *Wapl* KO conditions. Briefly, correlation matrices are derived from data by taking the pairwise linear correlation coefficient between each pair of columns in the Hi-C contact matrices. Then, a null hypothesis is defined from corresponding population correlation matrices. We then compute a Fisher z-transformation on the correlation matrices and calculate a test statistic for the chi-squared distribution to determine whether or not the null hypothesis (that Hi-C matrices are equivalent) should be rejected with a corresponding p-value.

### A/B Compartmentalization

Chromatin can either take on a condensed heterochromatic form or a looser euchromatic form. We use the Fiedler vector – the eigenvector corresponding to the second smallest eigenvalue (Fiedler number) of the normalized Laplacian matrix – to describe this feature mathematically which allows us to bi-partition the data into signed compartments – “A” (positive values) corresponding to euchromatin and “B” (negative values) corresponding to heterochromatin. The Fiedler vector and first principal component (PC1) vector are mathematically equivalent. However, unlike the PC1 vector, the Fiedler vector is accompanied by its corresponding Fiedler number which allows us to assign a magnitude of connectivity to a network. We characterize compartment switch events by identifying genomics bins with differing sign and magnitude above the 90th percentile of vector values between settings.

### Identification of TADs

Topologically associating domains (TADs) were identified using spectral clustering as described in chen *et al*. (Chen et al., 2016). This technique calculates the Fiedler vector for a normalized Hi-C adjacency matrix and initially organizes neighboring regions whose Fiedler vector values have the same sign into shared domains. This initial TAD structure is repartitioned if for a given domain, the Fiedler number is less than a user-defined threshold (thr) to ensure that the TADs represent well-connected regions and are not too large. Resulting TADs are iteratively repartitioned until their respective Fiedler numbers are larger than *λ_thr_* or until the smallest allowable TAD size (default is 300kb) is reached. We perform this procedure for chromosome-level Hi-C contact maps pooled across samples at 50kb resolution. For our analysis, thr was chosen to ensure a median TAD size of 900kb, as the expected median TAD size in mammalian genomes is 880kb (rounded to 900kb for our data since bins are in intervals of 50kb) (Dixon et al., 2012; Bonev and Cavalli, 2016). A specific *λ_thr_* was chosen for each chromosome and sample to ensure each TAD clustering set would have the same approximate median TAD size.

### Quantifying internal TAD organization

The internal structure of TADs is comprised of transient interactions over varying lengths that form chromatin ‘loops’. In the absence of WAPL, loop length increases, enriching contact frequency between either boundary of a given TAD (Haarhuis et al., 2017). Visually, we identify these interactions by the notable density of contacts present in the top right or bottom left corners of TADs. We quantify the strength of these interactions by designating a local neighborhood with a window size of 150kb by 150kb centered around the corner of each TAD. We then find the nonzero average of observed contacts in each window. TADs that are less than or equal to the window size are ignored (corner peak signal set to 0). This is performed for every TAD across each setting and chromosome. To evaluate the statistical significance of a given change in corner peak signal between WT and KO cells, we compared each corner peak signal to a background set of interactions. This background set is made up of contacts in the region between the diagonals encompassing the corner peak neighborhoods. We evaluate the p-value of each corner peak signal based on the proportion of background interactions with a signal less than the corner peak signal. As such, we determined that if a corner peak has a higher signal in the KO cells compared to WT cells and is significantly higher than interactions at comparable distances (p *<* 0.01), then the corner peak enrichment is statistically significant.

### Hi-C derived 5C contact map generation

We constructed a synthetic 5C contact map derived from a genome-wide 1Mb Hi-C contact map for genomic regions containing genes in the cell cycle gene networks. This was done by locating the genomic bins corresponding to cell cycle genes, extracting the inter-chromosomal and cell cycle loci-specific intra-chromosomal interaction frequencies for those bins, and stitching them together in genomic order. Our working set of cell cycle genes was sourced from the KEGG database and included 170 genes distributed across all chromosomes in the genome. Some of the genes occupy the same 1Mb genomic bins resulting in a 141 by 141 dimension adjacency matrix at 1Mb-resolution.

### Cell proliferation assay

CellTrace™Far Red was utilized for our cell proliferation assay. Purified T cells were labeled with far-red at a final concentration of 1 mol/L according to the manufacturer’s instruction (Molecular Probes, C34564). Far Red-labeled T cells were cultured similarly to above MLR for allogeneic reaction or Dynabeads T cell activator CD3/CD28 stimulation for up to 7 days. Then, collected cells were examined with flow cytometry, gated for CD3 (APC) positive, and Far Red dilution was determined.

### FxCycle™Far Red Stain for DNA content measurement

For *in vitro* experiments, WT or *Wapl* KO T cells were treated with Dynabeads T cell activator CD3/CD28 stimulation for up to 4 days. For *in vivo* experiments, transferred WT and *Wapl* KO T cells were isolated and purified from spleens of BALB/c recipient mice on day 7 after allogeneic BMT. The T cells were fixed with 10% formaldehyde for 30 mins, washed 3 times with PBS and the sample cell concentration was adjusted at 1 × 106 cells/mL. FxCycle™ Far Red stain (200 nM) (Molecular Probes, F10348) and 5 L of RNase A (20 mg/mL) (Roche Cat. 70294823) were added to flow cytometry samples and continued for incubation at room temperature for 30 minutes and protected from light. Samples were analyzed with flow cytometer (AttuneNxT) without washing using 633nm excitation and emission collected in a 660 bandpass. The DNA contents were determined as 2C and *>*2C by fluorescence intensities.

### ELISA

Concentrations of TNF-*α* and IFN-*γ* in sera on day 21 after allogeneic BMT were measured with specific anti-mouse ELISA kits from BD Biosciences. Assays were performed per the manufacturer’s protocol and read at 450nm in a microplate reader (Bio-Rad). The concentrations were calculated from triplicate samples as mean ± SEM.

### FACS

Single-cell suspensions of spleens and thymii were prepared as previously described (Sun et al., 2015). Briefly, to analyze surface phenotype, purified T cells and thymocytes from B6 WT, *Wapl* KO deficiency mice, or transplanted animals, were washed with FACS wash buffer (2% bovine serum albumin [BSA] in phosphate-buffered saline [PBS]), pre-incubated with the rat anti-mouse FcR mAb 2.4G2 for 15 minutes at 4°C to block nonspecific FcR binding of labeled antibodies, then resuspended in FACS wash buffer and stained with conjugated monoclonal antibodies purchased from BD Biosciences (San Jose, CA): allophycocyanin (APC)-conjugated monoclonal antibodies (MoAbs) to CD4, CD8, CD3, CD45.2, CD45.1, CD25 and CD69; phycoerythrin (PE)-conjugated MoAbs to CD3, CD4, CD8, CD25, and TCR*β*; allophycocyanin (APC)-conjugated MoAbs to CD3, CD4, and PerCP/Cy5.5-conjugated MoAbs to CD3, CD4 and CD8 were purchased from eBioscience (SanDiego, CA). Next, cells were analyzed using an AttuneNxT flow cytometer. For intracellular staining, cells were stained for CD4 and CD25 antibodies as above, then fixed with IC Fixation Buffer (Biolegend, Cat. No.420801) and incubated 20-60 minutes at room temperature and continued by adding 2 mL of 1X Permeabilization Buffer (Biolegend Cat. No. 421002) and centrifuged at 400-600 x g for 5 minutes at room temperature. Cell pellets were resuspended in 100 L of 1X permeabilization buffer and PE-conjugated FoxP3 antibody (eBioscience, SanDiego, CA, Cat. 126403) was added at 0.5 ug/million cells/100 ul and incubated for 30 minutes at room temperature. Stained cells were resuspended in an appropriate volume of Flow Cytometry Staining Buffer for flow cytometry analyses. Apoptotic cells were detected by PE-Annexin V staining.

---

**Fig. S1:**
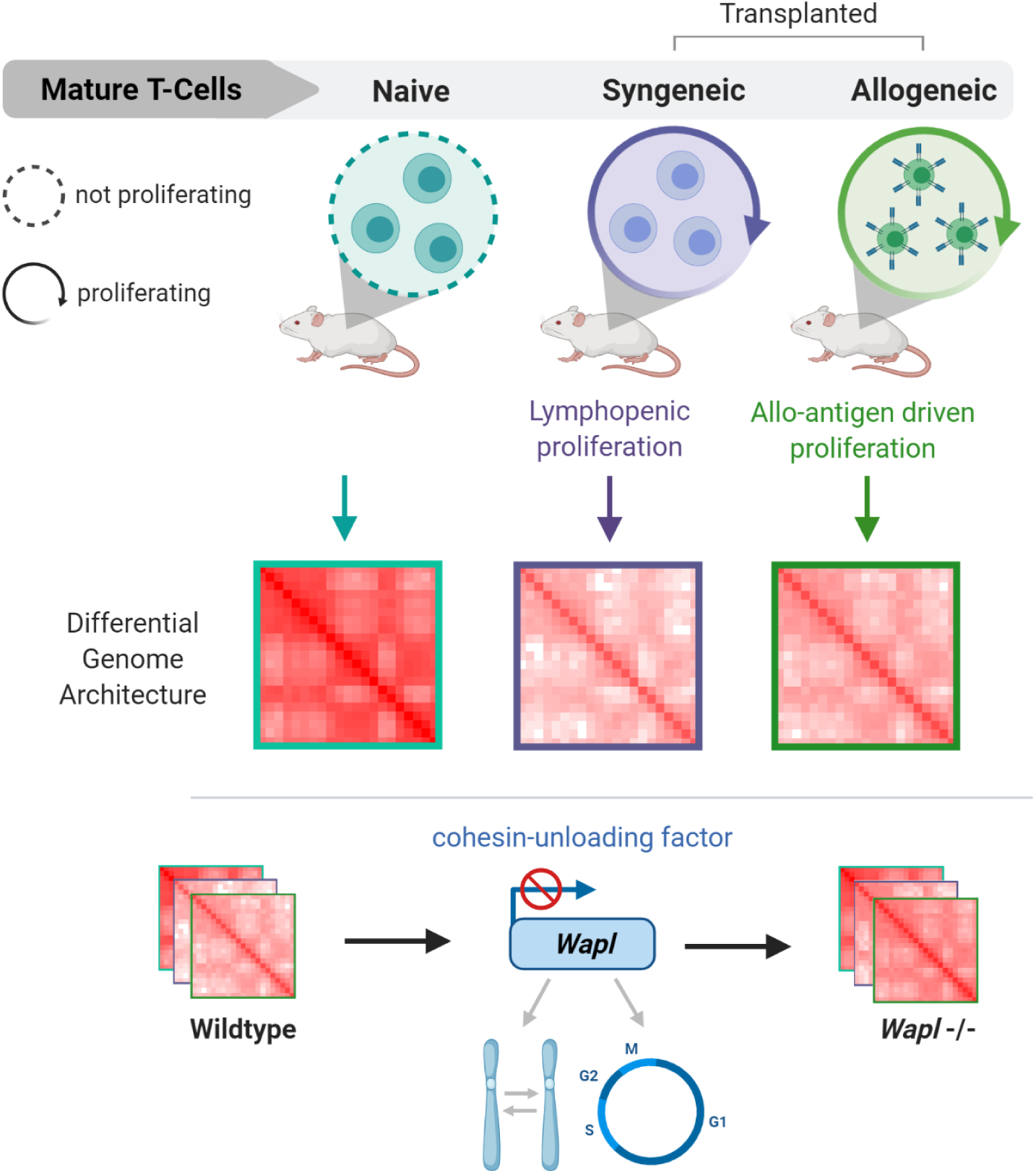
Overview, related to STAR Methods. We combined genome-wide chromosome conformation capture (Hi-C) with RNA-sequencing to describe the 3D chromatin architecture of mature T cells *in vivo*, at baseline (naive) and following lymphopenic proliferation (syngeneic) and allo-antigen driven (allogeneic) proliferation. We generated T cell-specific WAPL-deficient mice to explore whether WAPL regulates the 3D chromatin structure, function, and *in vivo* biological response of T cells.

**Fig. S2:**
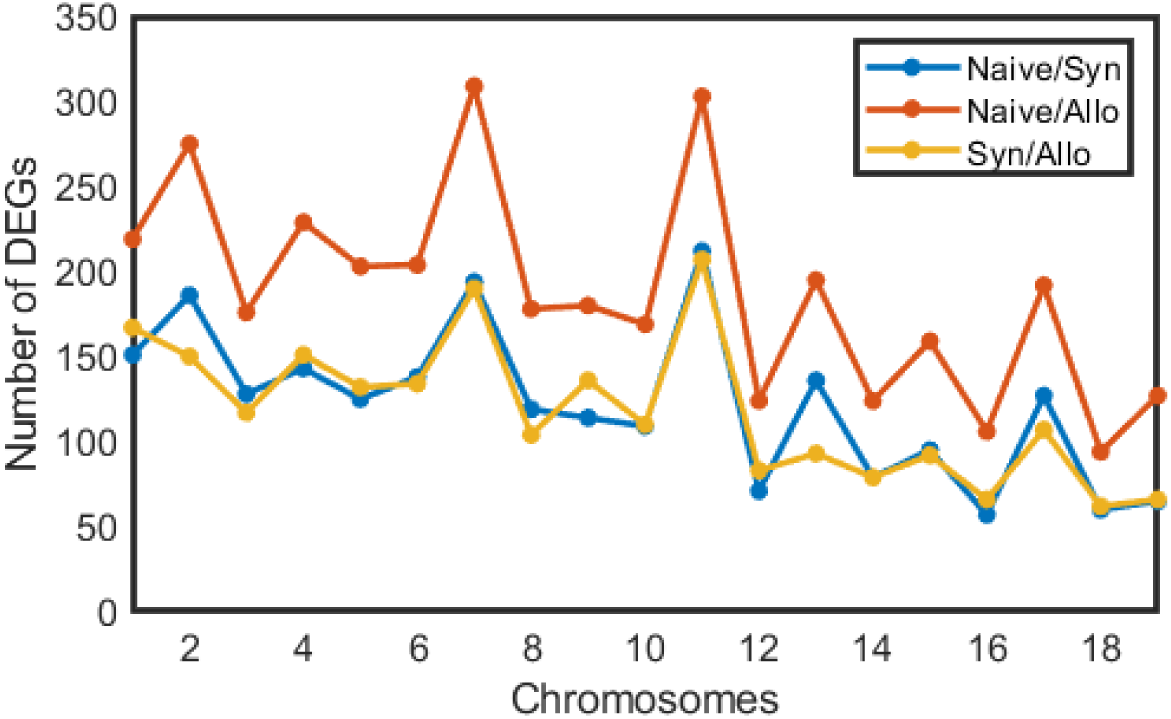
Differential Expression Between Unstimulated and Stimulated T Cells, related to Figure 2. A total of 2,309 genes were significantly differentially expressed between wildtype naive and syngeneic T cells across all chromosomes (log_2_FC≥1, p≤0.001). Significant differential expression was identified for 3,566 genes between wildtype naive and syngeneic T cells and 2,246 genes between wildtype syngeneic and allogeneic T cells.

**Fig. S3:**
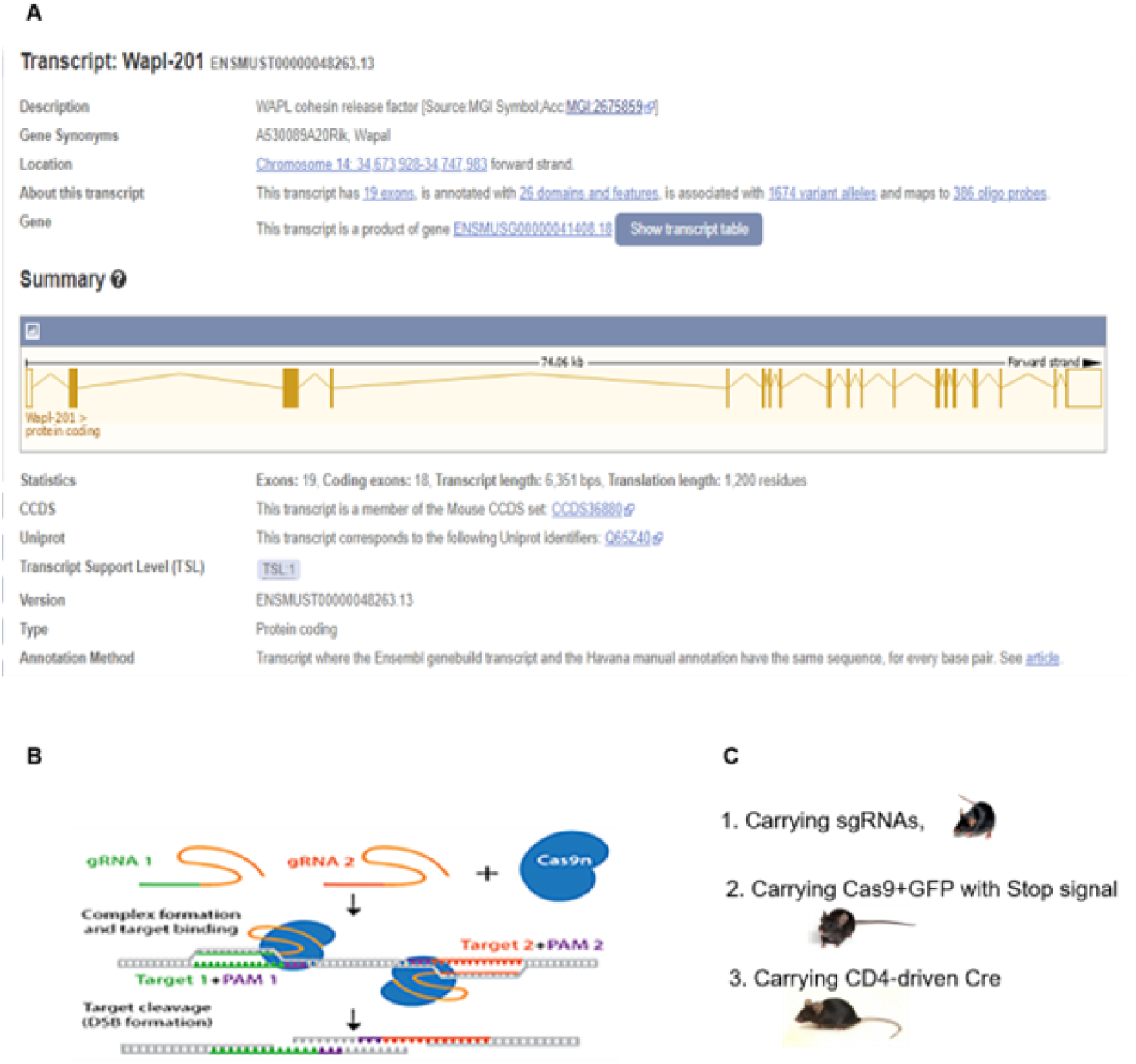
*Wapl* gene locus and targeting strategy for sgRNAs-CRISPR-CAS9, related to Figure 1. (A) The *Wapl* locus is localized in mouse Chromosome 14 qB with 18 coding exons and 1 noncoding exon. (B) Mechanism of production of double string break (DSB) in exon 2 of *Wapl* gene by 2 sgRNAs-CRISPR-CAS9 system. (C) 3 mice lines with B6 background used in generation of conditional *Wapl* KO in T cells: 1) Generated mouse which carries an insert of 2 sgRNAs targeting exon 2 in *Wapl* gene; 2) Rosa26-floxed STOP-Cas9^+^GFP knockin mouse (Rosa26-LSL-Cas9 knockin on B6J) which carries Cas9 and GFP with stop signal; 3) CD4CRE which carries CD4-driven CRE recombinase.

**Fig. S4:**
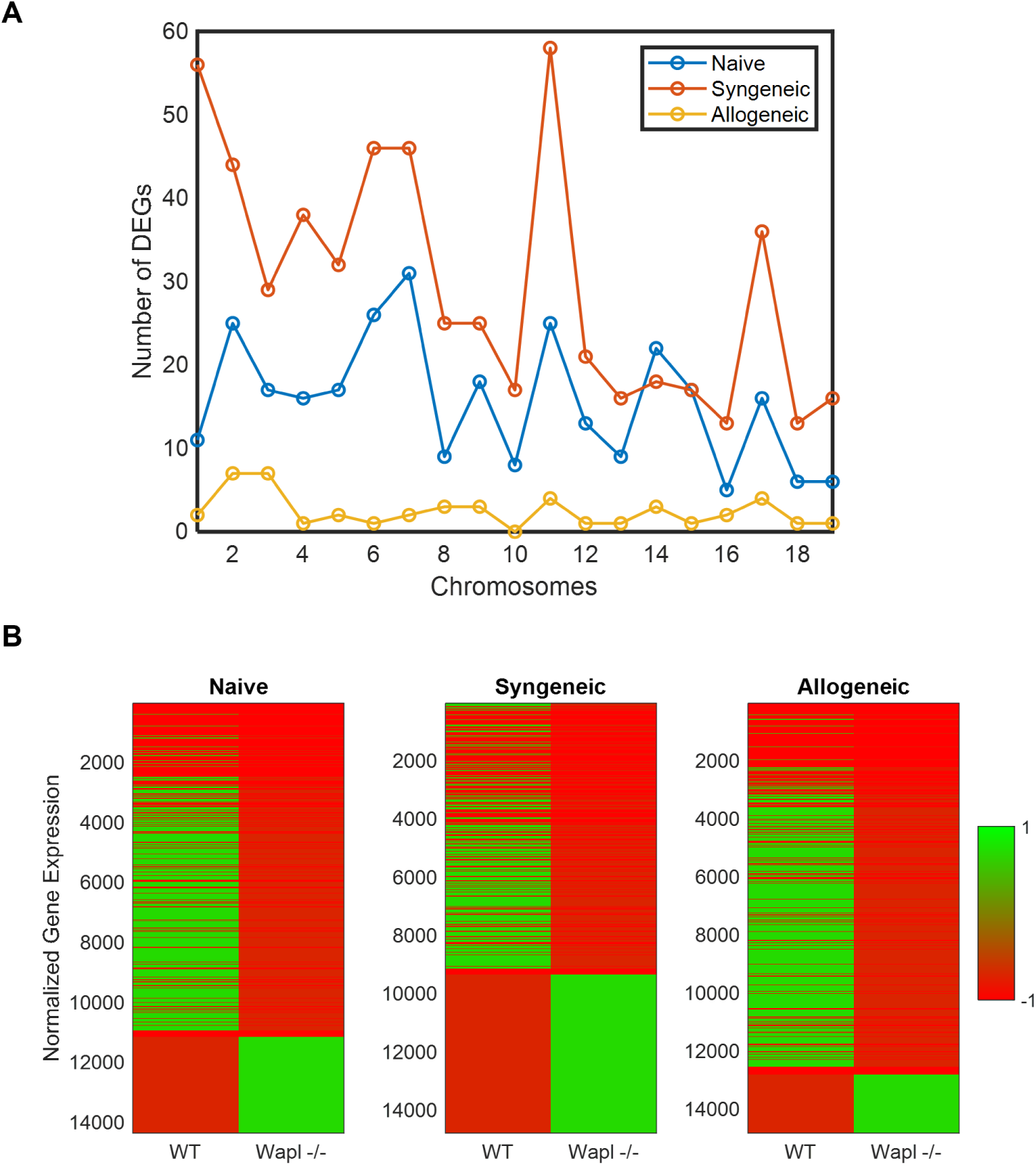
*Wapl* Knockout Differential Expression Summary, related to Figure 2. (A) Number of significantly differentially expressed genes (log_2_FC≥1, p≤0.01) between WT and KO per chromosome and setting. (B) Differential expression clusters across all genes.

**Fig. S5:**
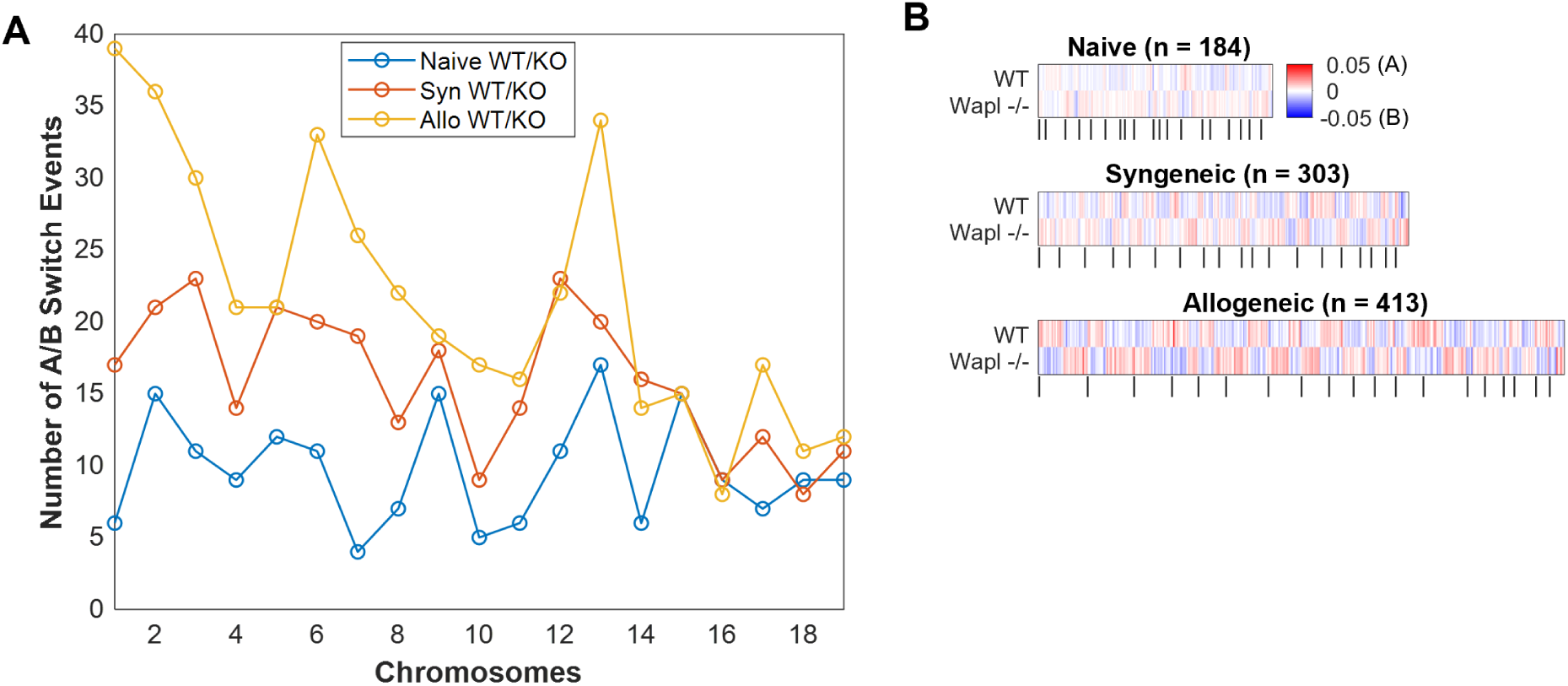
A/B compartment summary, related to Figure 2. (A) Distribution of number of A/B compartment switch events per chromosome between WT and KO settings. Each setting is a different color. (B) A/B compartment switch events between WT and KO T cells. We observed 184 loci (100kb-length) with different chromatin states between WT and KO unstimulated naïve T cells, 303 between both WT and KO syngeneic T cells, and 413 between WT and KO allogeneic T cells. Vertical bars below each heatmap indicate chromosomes in numerical order (1 to 19) from left to right. The heatmap color intensity reflects the magnitude of the Fiedler values for each locus.

**Fig. S6:**
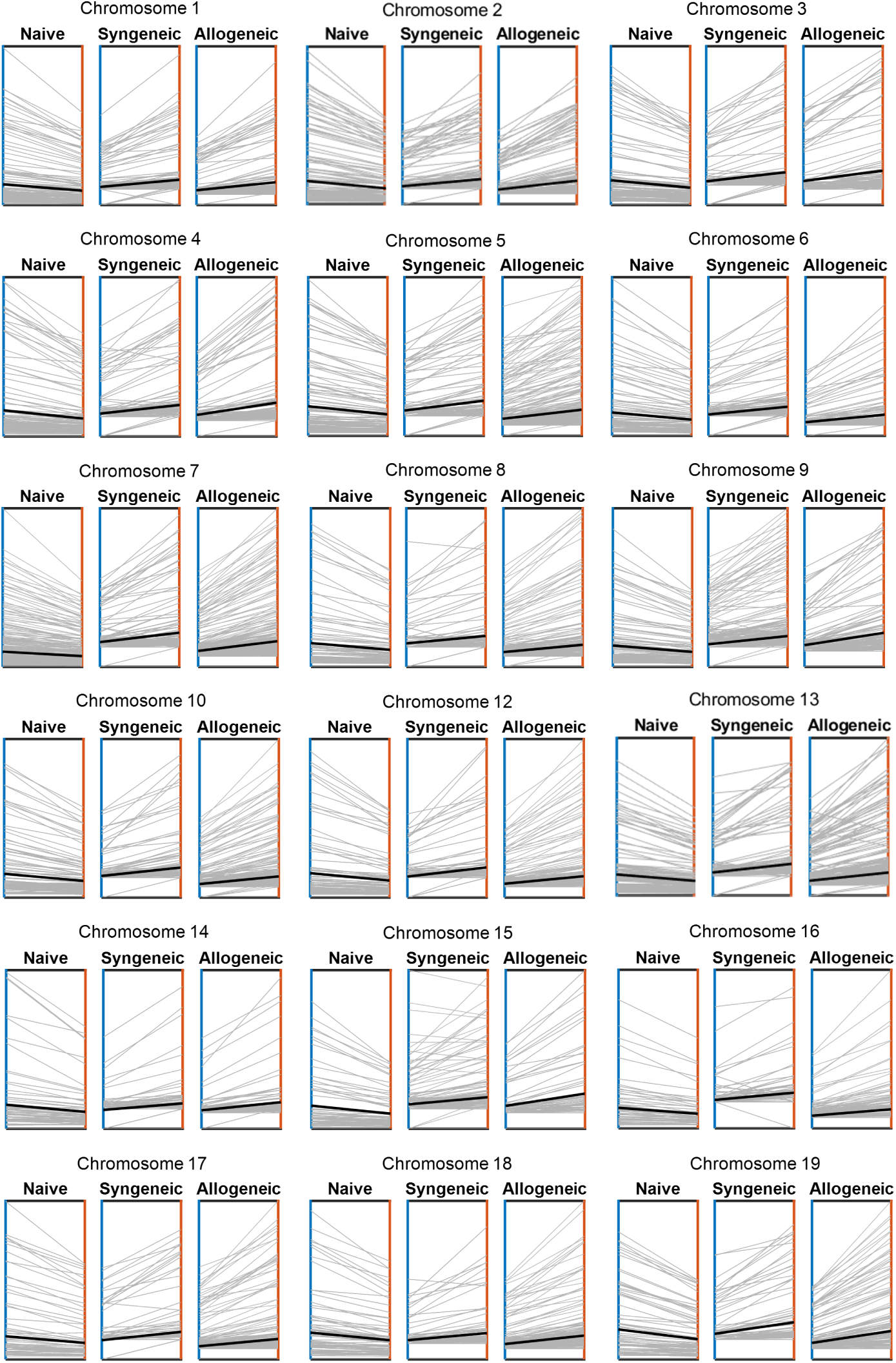
Corner Peak Dynamics, related to Figure 3. Change in corner peak signal from WT (blue axis) to KO (orange axis) for each TAD (individual gray lines). The average change in corner peak signal between the two conditions is plotted as a solid black line in each panel to capture the overall trend across settings. TADs and their corner peak signals are represented at 50kb resolution. Chromosome 11 is omitted from this figure as it is already highlighted in Figure 3

**Fig. S7:**
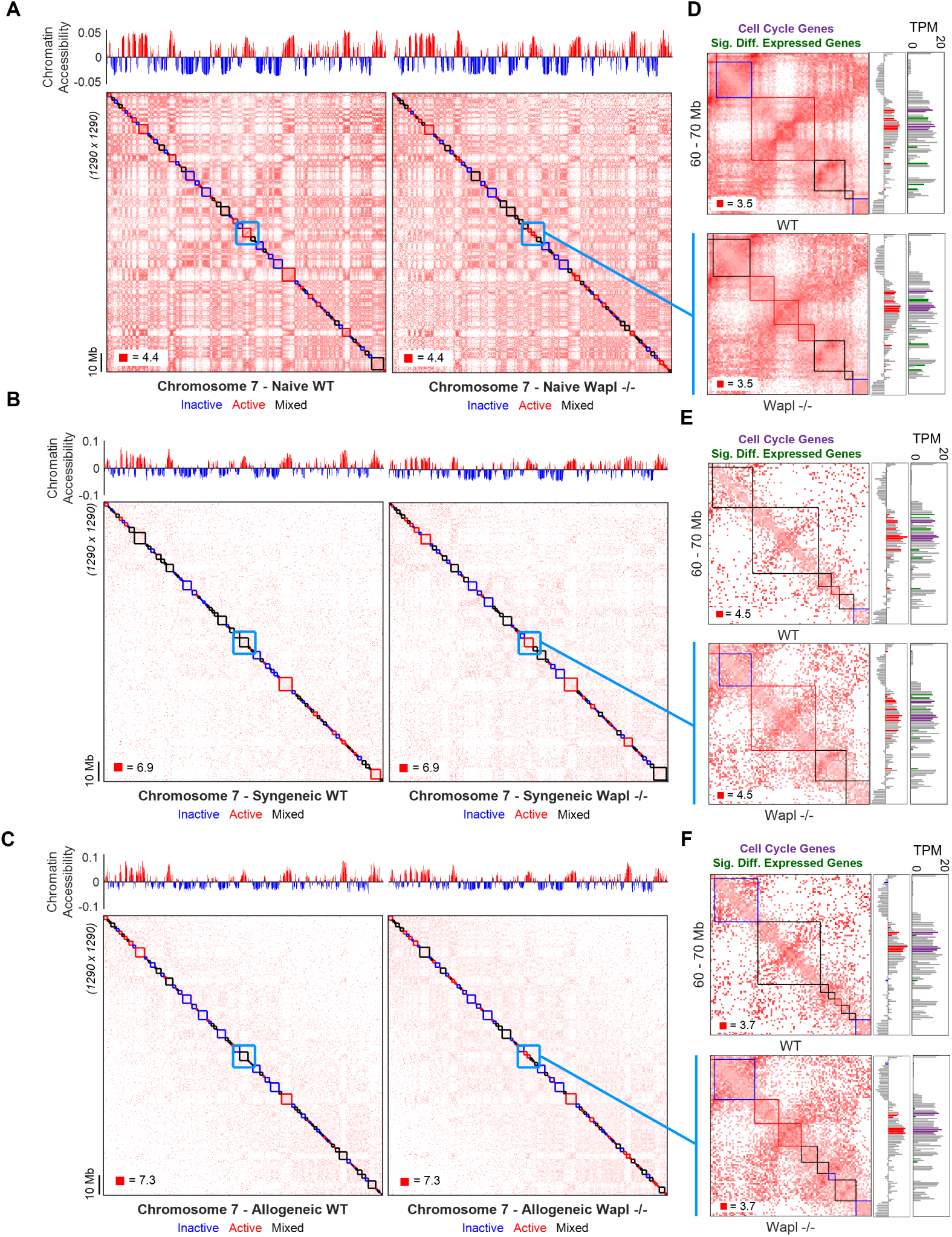
Comparison of TAD organization between WT and KO T cells, related to Figure 3. (A-C) Chromosome 7 comparison of chromatin accessibility (top, track) and TAD clustering (bottom, map) between WT and KO unstimulated naïve (A), syngeneic (B), and allogeneic (C) T cells. TADs are shown along a normalized (observed/expected) log-scale contact map at 100-kb resolution and are colored according to chromatin accessibility with TADs containing both accessible and inaccessible loci denoted ‘mixed’ and colored black. (D-F) The light blue boxed area on the contact maps highlights gene-rich region 60Mb – 70Mb in Chromosome 7. Zoomed in panels to the right offer a closer look at the region’s interaction pattern and TADs, chromatin accessibility (bar plot directly right), and gene expression (rightmost bar plot). This region contains three cell cycle genes (*Fanci*, *Prc1*, *Blm*), whose chromatin accessibility is indicated by red bars in the chromatin accessibility plot and whose gene expression is indicated by the purple bars in the gene expression plot. There are more than three highlighted bars in both plots because the three genes each spanned two 100kb-length regions/bins. Green bars in the gene expression plot represent 100kb-length regions/bins containing one or more non-cell cycle genes that are differentially expressed (*|*log_2_FC*| ≥* 1).

**Fig. S8:**
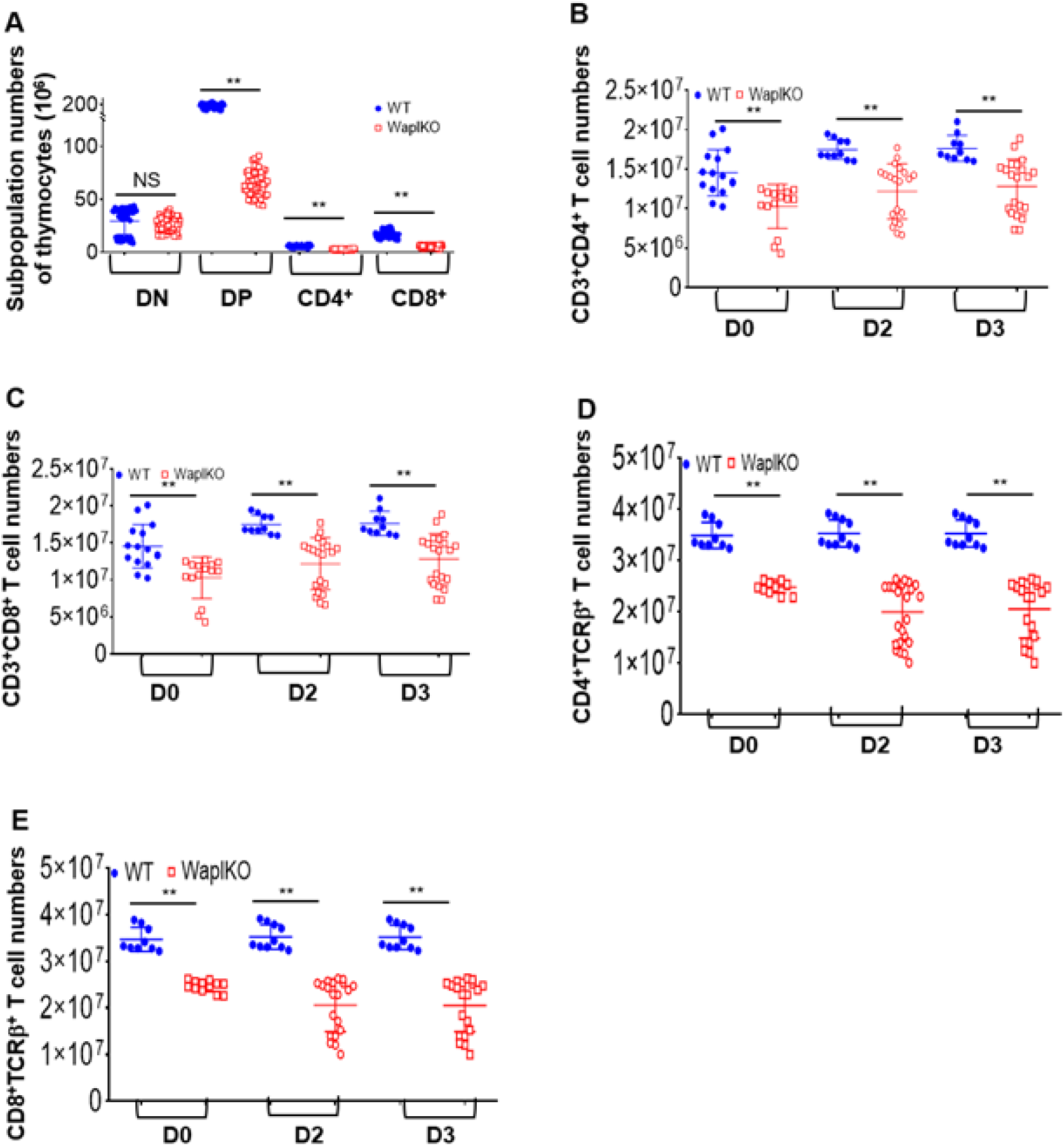
WAPL deficiency impairs T cell development and function, related to Figure 5. (A) WT and *Wapl* KO thymocytes were isolated and stained with CD4 and CD8 antibodies and analyzed by flow cytometry. The numbers of thymocytes for double negative (DN), double positive (DP), CD4 or CD8 single positive were calculated. Data were combined from thymuses of 9 WT and 10 *Wapl* KO mice (mean ± SEM). (B-E) Purified WT and *Wapl* KO T cells were left in a naïve state or stimulated with CD3/CD28Ab for up to 3 days, stained with CD3, CD4, CD8, TCR antibodies, and analyzed by flow cytometry. The T cell numbers either WT or *Wapl* KO for CD3^+^CD4^+^(B), CD3^+^CD8^+^ (C), CD4^+^TCR^+^ (D) and CD8^+^TCR^+^ (E) were calculated. Data were combined from 3 independent experiments (mean ± SEM).

**Fig. S9:**
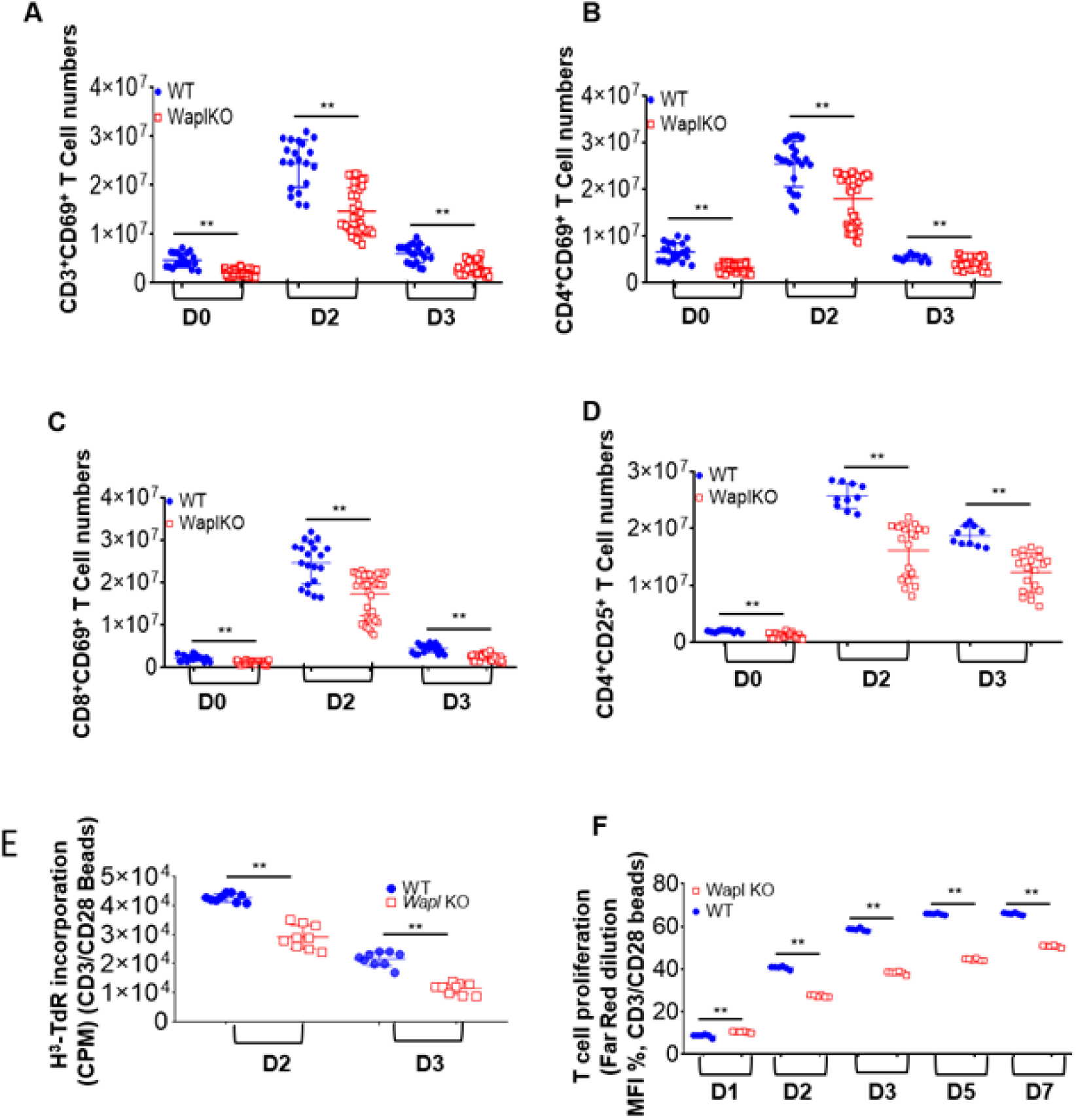
WAPL deficiency impairs T cell function, related to Figure 5. (A-D) Purified WT and WAPL-deficient T cells were left untreated in a naïve state or stimulated with CD3/CD28Ab for up to 3 days, then harvested for FACS staining. The T cell numbers either WT or *Wapl* KO for CD3^+^CD69^+^ (A), CD4^+^CD69^+^ (B), CD8^+^CD69^+^(C), and CD4^+^CD25^+^ (D) were analyzed and calculated. Data were combined from 3 independent experiments (Mean ± SEM). (E) H3-TdR incorporation in WT and *wapl* KO T cells were assessed by mixed lymphocyte reaction when stimulated with CD3/CD28mAb for up to 3 days. Data were combined from 2 independent experiments (mean ± SEM). (F) Cell proliferation was analyzed by Far Red dilution for either WT or *Wapl* KO T cells when stimulated with CD3/CD28Ab for up 7 days. Data were combined from 2 independent experiments (mean ± SEM).

**Fig. S10:**
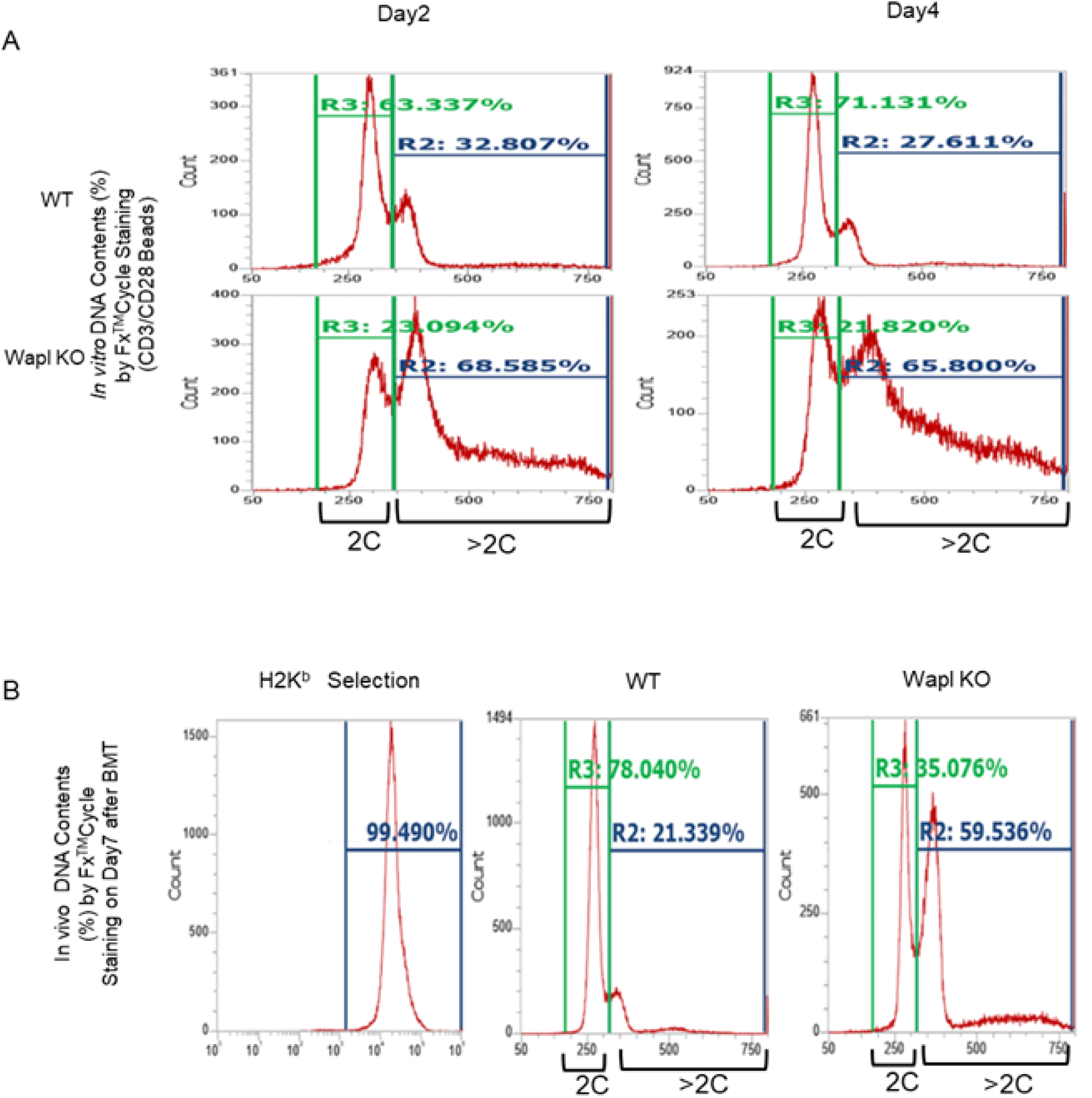
WAPL deficiency in T cells impairs T cell cycling *in vitro* and *in vivo*, related to Figure 5. (A) After WT and *Wapl* KO T cells were stimulated with CD3/CD28AB for 2-4 days *in vitro*, cells were stained with FxCycle™Far Red, DNA contents were separated as 2C and *>*2C by flow cytometry. A representative assay is shown. (B) WT and *Wapl* KO T cells were isolated from recipient spleens on day 7 after allogeneic BMT *in vivo* and stained with FxCycle™Far Red. DNA contents were separated as 2C and *>*2C by flow cytometry. A representative assay is shown.

**Table S1:**
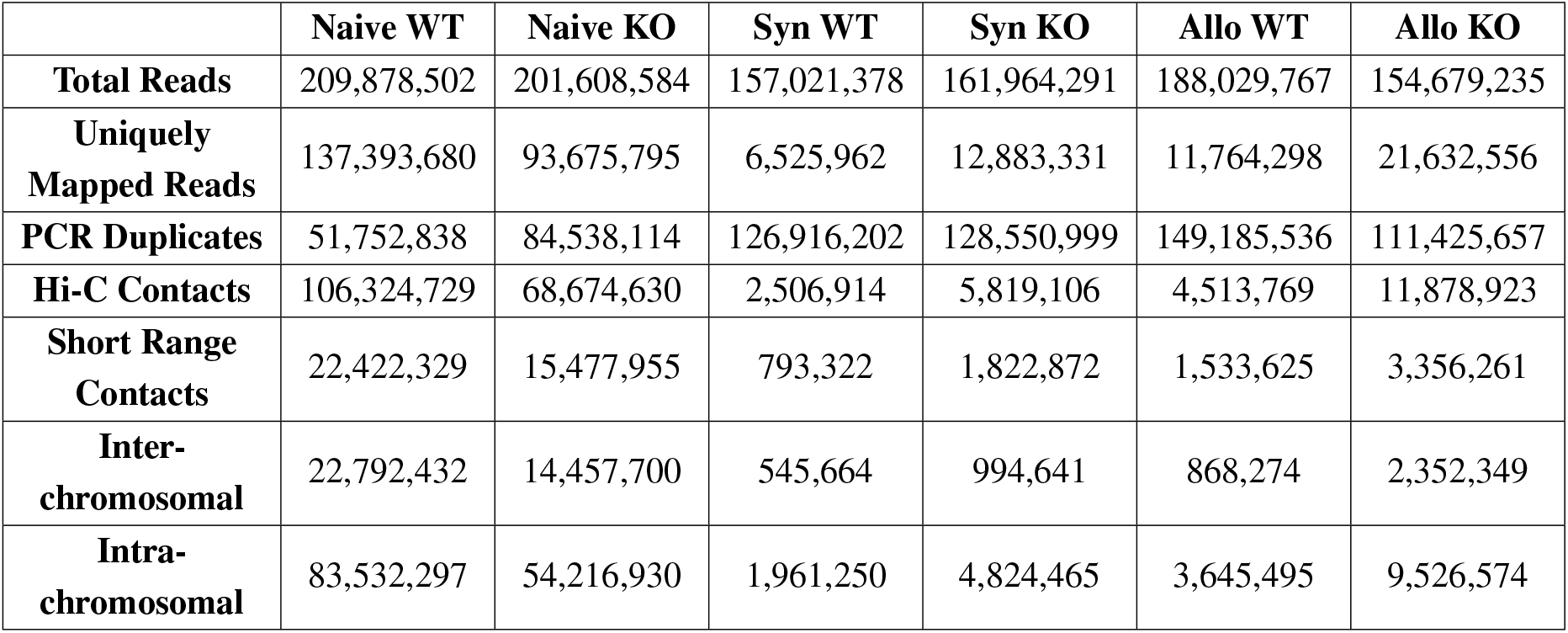
Hi-C QC Summary.

|  | Naive WT | Naive KO | Syn WT | Syn KO | Allo WT | Allo KO |
| --- | --- | --- | --- | --- | --- | --- |
| **Total Reads** | 209,878,502 | 201,608,584 | 157,021,378 | 161,964,291 | 188,029,767 | 154,679,235 |
| **Uniquely Mapped Reads** | 137,393,680 | 93,675,795 | 6,525,962 | 12,883,331 | 11,764,298 | 21,632,556 |
| **PCR Duplicates** | 51,752,838 | 84,538,114 | 126,916,202 | 128,550,999 | 149,185,536 | 111,425,657 |
| **Hi-C Contacts** | 106,324,729 | 68,674,630 | 2,506,914 | 5,819,106 | 4,513,769 | 11,878,923 |
| **Short Range Contacts** | 22,422,329 | 15,477,955 | 793,322 | 1,822,872 | 1,533,625 | 3,356,261 |
| **Inter-chromosomal** | 22,792,432 | 14,457,700 | 545,664 | 994,641 | 868,274 | 2,352,349 |
| **Intra-chromosomal** | 83,532,297 | 54,216,930 | 1,961,250 | 4,824,465 | 3,645,495 | 9,526,574 |

**Table S2:** Summary of Chromatin Accessibility Changes After *Wapl* Knockout.

| | Stable A (%) | Stable B (%) | A → B (%) | B → A (%) |
| --- | --- | --- | --- | --- |
| **Naive** | 49.8 | 49.4 | 0.2 | 0.6 |
| **Syngeneic** | 49.9 | 48.8 | 0.5 | 0.8 |
| **Allogeneic** | 46.8 | 51.4 | 0.8 | 0.9 |

**Table S3:** Percent of TADs With Statistically Significant Corner Peak Signal (p *<* 0.01)

|  | Naive |  | Syngeneic |  | Allogeneic |  |
| --- | --- | --- | --- | --- | --- | --- |
| Chr | WT | KO | WT | KO | WT | KO |
| 1 | 62% | 64% | 34% | 41% | 31% | 46% |
| 2 | 56% | 58% | 31% | 34% | 37% | 45% |
| 3 | 66% | 69% | 31% | 42% | 37% | 49% |
| 4 | 60% | 63% | 33% | 35% | 38% | 45% |
| 5 | 62% | 65% | 33% | 37% | 37% | 49% |
| 6 | 55% | 61% | 20% | 31% | 32% | 42% |
| 7 | 64% | 65% | 32% | 34% | 46% | 55% |
| 8 | 54% | 61% | 25% | 31% | 37% | 42% |
| 9 | 58% | 65% | 28% | 28% | 38% | 48% |
| 10 | 59% | 66% | 30% | 35% | 35% | 47% |
| 11 | 52% | 57% | 24% | 26% | 32% | 43% |
| 12 | 58% | 62% | 27% | 33% | 28% | 38% |
| 13 | 64% | 64% | 35% | 32% | 39% | 43% |
| 14 | 52% | 54% | 25% | 23% | 27% | 32% |
| 15 | 56% | 59% | 36% | 32% | 29% | 40% |
| 16 | 56% | 62% | 22% | 25% | 26% | 37% |
| 17 | 44% | 49% | 25% | 24% | 30% | 38% |
| 18 | 66% | 68% | 26% | 33% | 30% | 41% |
| 19 | 58% | 64% | 30% | 38% | 33% | 44% |

**Table S4:** Differentially Expressed Cell Cycle Genes Between WT and KO T-Cells Across Settings (p *<* 0.001).

|  | Chromosome | Gene | log<sub>2</sub>(Fold Change) |
| --- | --- | --- | --- |
| **Naive** | 8 | *Mtus1* | 1.04 |
|  | 11 | *Gas2l1* | 1.12 |
|  | 6 | *Rassf4* | 1.21 |
|  | 17 | *Cdkn1a* | 2.20 |
|  | 14 | *Ajuba* | 2.71 |
| **Syngeneic** | 5 | *Evi5* | -3.09 |
|  | 2 | *Gfi1b* | -2.79 |
|  | 19 | *Anxa1* | -2.77 |
|  | 10 | *Sash1* | -2.38 |
|  | 9 | *Tfdp2* | -2.26 |
|  | 17 | *Aif1* | -2.15 |
|  | 11 | *Myh10* | -2.08 |
|  | 6 | *Asns* | -1.66 |
|  | 11 | *Sphk1* | -1.57 |
|  | 1 | *Steap3* | -1.20 |
|  | 8 | *Mtus1* | -1.10 |
|  | 8 | *Cul4a* | -1.04 |
|  | 3 | *Mnd1* | 1.10 |
|  | 14 | *Ajuba* | 1.39 |
|  | 13 | *Esm1* | 1.43 |
|  | 15 | *Krt18* | 1.46 |
| **Allogeneic** | 19 | *Anxa1* | -1.02 |

